# A cytochrome P450 G subfamily member, CYP4G35, is highly expressed in antennae and modulates olfactory response in *Aedes aegypti* mosquitoes

**DOI:** 10.1101/2025.11.13.688110

**Authors:** Arvind Sharma, Andrew Nuss, Omar Garcia-Cruz, Fiza Arshad, Jeremiah Reyes, Manoj Mathew, Marina MacLean, Sharon Young, Runzhi Hu, Saransh Beniwal, Claus Tittiger, Gary Blomquist, Michael Pham, Juli Petereit, Dennis Mathew, Robert Harrell, Monika Gulia-Nuss

## Abstract

The cytochrome P450 enzymes of the CYP4G subfamily are some of the most enigmatic insect P450s. The dipterans with sequenced genomes have two CYP4G paralogs. In *Drosophila melanogaster*, CYP4G1 is highly expressed in the oenocytes and catalyzes the last enzymatic step in the biosynthesis of cuticular hydrocarbons. In contrast, CYP4G15 is expressed in the brain glial cells, but its function is unknown. The *Aedes aegypti* genome encodes two CYP4Gs: CYP4G36 (ortholog of DmCYP4G1) and CYP4G35 (ortholog of DmCYP4G15). Here, we show that CYP4G35 is highly expressed in mosquito antennae, and the RNAi knockdown of CYP4G35 results in delayed host-seeking. *Ae. aegypti* CYP4G knockout lines confirmed delayed host-seeking behavior in CYP4G35 knockout females. Proteomics analysis of CYP4G35 KO females also corroborates the physiological findings and shows upregulation of proteins related to olfaction and other CYP4Gs to compensate for the lack of CYP4G35. Immunohistochemistry and *in situ* hybridization were used to localize CYP4G35 and demonstrated its expression in the sensilla lymph of the antennae and the tip of the proboscis. CYP4G35 and CYP4G36 fusion proteins with cytochrome P450 reductase demonstrated that, unlike CYP4G36, CYP4G35 lacks an oxidative decarbonylase function. Together, our data support a novel function of CYP4G35 in modulating olfactory response.

## INTRODUCTION

*Aedes aegypti* mosquitoes transmit arboviruses, including the causative agents of dengue, Zika, yellow fever, and chikungunya. Female *Ae. aegypti* mosquitoes require a blood meal for reproduction^1^ and prefer biting humans, which contributes to their effectiveness as disease vectors^2–4^. In addition to finding a vertebrate host for blood feeding and a suitable oviposition site, both females and males also need to find nectar sources. Consequently, mosquitoes, including *Ae. aegypti*, have evolved an acute olfactory system. The olfaction process begins with the reception of odor molecules by specialized peripheral sensory structures, known as olfactory sensilla, which are present on various chemosensory tissues, including antennae, maxillary palps, and proboscis. This is followed by the integration of olfactory and other sensory modalities in the brain, and ultimately, the translation of olfactory signals into behavior^5,6^. Thus, the cornerstone of a sophisticated olfactory system is the ability of the insect’s peripheral system to selectively detect and rapidly inactivate odorants once they have conveyed information.

*Aedes aegypti*, like other insects, detects chemosensory cues using receptors encoded by three large multi-gene families, odorant receptors (ORs), ionotropic receptors (IRs), and gustatory receptors (GRs). It has been hypothesized that odorant-degrading enzymes (ODEs) are likely involved in terminating odorant signals. A global scheme has been proposed to explain most molecular interactions, such as the transport of odor molecules by odorant-binding proteins (OBPs) through the sensillum lymph and their interaction with chemosensory receptors^6–9^. Odorant molecules are rapidly removed to allow detection of new stimuli, most likely by extracellular ODEs.

To satisfy the criteria for an ODE, an enzyme must 1) reside in the sensilla lymph and 2) degrade an odorant/ volatile compound. These criteria have been fulfilled by antennal esterases, which degrade pheromones and are classified as pheromone-degrading enzymes (PDEs)^6^. *In vitro* experiments have also demonstrated that antennal aldehyde oxidases, aldehyde dehydrogenases, epoxide hydrolases, glutathione-S-transferases, and cytochrome P450s degrade various pheromones^6^.

Cytochromes P450 (CYP) enzymes constitute one of the largest gene families among all living organisms. Over 100 CYPs have been identified in the *Ae. aegypti* genome^10^. CYPs are heme thiolate proteins known to catalyze xenobiotic metabolism and epoxidation. The insect CYPs can be divided into four major clans: CYP2, CYP3, CYP4, and mitochondrial CYPs^11^. Among CYP4s, the CYP4G subfamily is evolutionarily conserved across insects^12,13^ but is absent in other classes, such as Crustacea and Chelicerata^14^. An extra N-terminal loop is a characteristic feature of CYP4G subfamily enzymes, distinguishing them from other CYPs^12^. Dipteran genomes sequenced to date contain one or two CYP4Gs.

The *Drosophila melanogaster* genome has two CYP4G genes: CYP4G1 and CYP4G15^15^. CYP4G1 is the most highly expressed CYP and is massively co-produced with NADPH cytochrome P450 reductase (CPR) in oenocytes^16^. All of the active CYP4Gs assayed to date are oxidative decarbonylases^12,13,17–19^

In addition to CHC production, CYP4Gs also appear to have functions in phytochemical/ olfactory cue clearance. For instance, honeybee, *Apis, mellifera*, CYP4G11, the only CYP4G present in the honeybee, is highly expressed in antennae and legs^20^. CYP4G11 can accept short-chain aldehydes as substrates^13,21^ and convert these into volatile hydrocarbons that subsequently dissipate from the insect. Therefore, in addition to CHC production, these enzymes might be involved in clearing shorter-chain pheromonal and phytochemical alcohols and aldehydes by converting them to more volatile hydrocarbons^13^. In the armyworm, *Mamestra brassicae*, CYP4G20 transcripts were abundant in the antennae of adult males and were also present in all chemosensory tissues tested, including the male proboscis, legs, and female ovipositors, which possess taste sensilla, suggesting a function in olfaction and taste^22^. The *D. melanogaster* CYP4G15 is expressed in the nervous system^23^ and was recently shown to localize in the cortex glial cells in the brain^16^. However, its function is still unknown.

Our localization data meet criterion 1 for an ODE, and the behavioral data are consistent with an olfactory role. While we tested oxidative decarbonylation, we did not assess the metabolism of human host-relevant odorants by CYP4G35. Therefore, direct biochemical evidence for odorant degradation by CYP4G35 remains to be established. Together, our data are consistent with a role for CYP4G35 in modulating olfactory response, and future work defining its odorant substrates will complete the biochemical criteria for ODE function.

## Materials and Methods

### Mosquito Rearing

*Aedes aegypti* (LVP strain) colonies were maintained at 27 °C and 60-70% relative humidity (RH) under a 16:8 h light: dark photoperiod in a dedicated insectary. Eggs were hatched overnight in small plastic cups in deionized water. First instar larvae were counted (100 per 1.5-gallon pan) and reared in 500 mL of deionized water on a powdered fish food diet (Tetramin®, Melle, Germany), as described in detail in^24^. Pupae were collected from the rearing container and transferred to adult emergence cages. Adults were fed a 10% sucrose solution containing food color for regular colony maintenance. Adult females were blood-fed four days after emergence, and egg cups lined with a paper towel were placed in the cage 48 h after blood feeding. The egg cups were removed from the cages 3 days later; egg sheets were washed and dried overnight at room temperature (RT) and stored in a plastic bag.

### RNA extraction and cDNA synthesis

Expression of CYP4G35 (NCBI reference number: XM_001658018; also referred to as CYP4G15) and CYP4G36 (NCBI reference number: XM_001648326.2, also referred to as CYP4G15) was examined using qRT-PCR. Total RNA was extracted from eggs, 2^nd^ and 4^th^ instar larvae, early and late pupae, sugar and blood-fed females (24h post blood feeding), and sugar-fed males and female tissues (head, antennae, proboscis, maxillary palps, midgut, ovaries, fat body/abdomen, and pro-, meso-, and metathoracic legs). A pool of 5-20 mosquitoes, depending on the stage (three technical and three biological replicates) was used for RNA isolation using TRIzol reagent according to the manufacturer’s protocols (Invitrogen, Waltham, MA, USA). RNA quantity was measured with a Nanodrop spectrophotometer. For adult female tissues, 50 tissues were pooled per replicate.

5 μg of total RNA was used for DNase treatment (Sigma, St. Louis, MO, USA) according to the manufacturer’s protocol. DNase-treated RNA samples were re-purified with Trizol. Purity was determined by the 260/280 and 260/230 ratios, with acceptable values in the 2.0 and 2.0–2.2 ranges, respectively (all samples collected met the purity standards). 1 μg of DNase-treated RNA was used for cDNA synthesis with iScript reverse transcription supermix (BioRad, Hercules, CA, USA). The cDNA was diluted 10-fold in water before being used as a template in qRT-PCR experiments.

### Quantitative RT-PCR

1 μl cDNA was used in each 10 μl qRT-PCR reaction. Each sample was run in quadruplet wells of a 96-well plate. qRT-PCR was performed on a CFX Touch Real-Time PCR Detection System (Bio-Rad, Hercules, CA, USA) using SYBR Green Master Mix (Bio-Rad). CYP4G specific primers (Table 1). A ribosomal protein S7 gene was used as a housekeeping control. All reactions were performed for 3 min at 95 °C, followed by 39 cycles of 10 s at 95 °C, 30 s at 58 °C, and 30 s at 72 °C, and the melt curve was analyzed at 65–95 °C. Relative expression was calculated using the 2^−ΔΔCt^ method^25^, where eggs were considered the control, and expression of CYP4G35 and CYP4G36 in other life stages was calculated relative to the control levels. For tissues, the ovaries were used as a control due to their very low expression in the preliminary experiments. Each experiment consisted of four technical and three biological replicates from different mosquito cohorts.

**Table 1:**
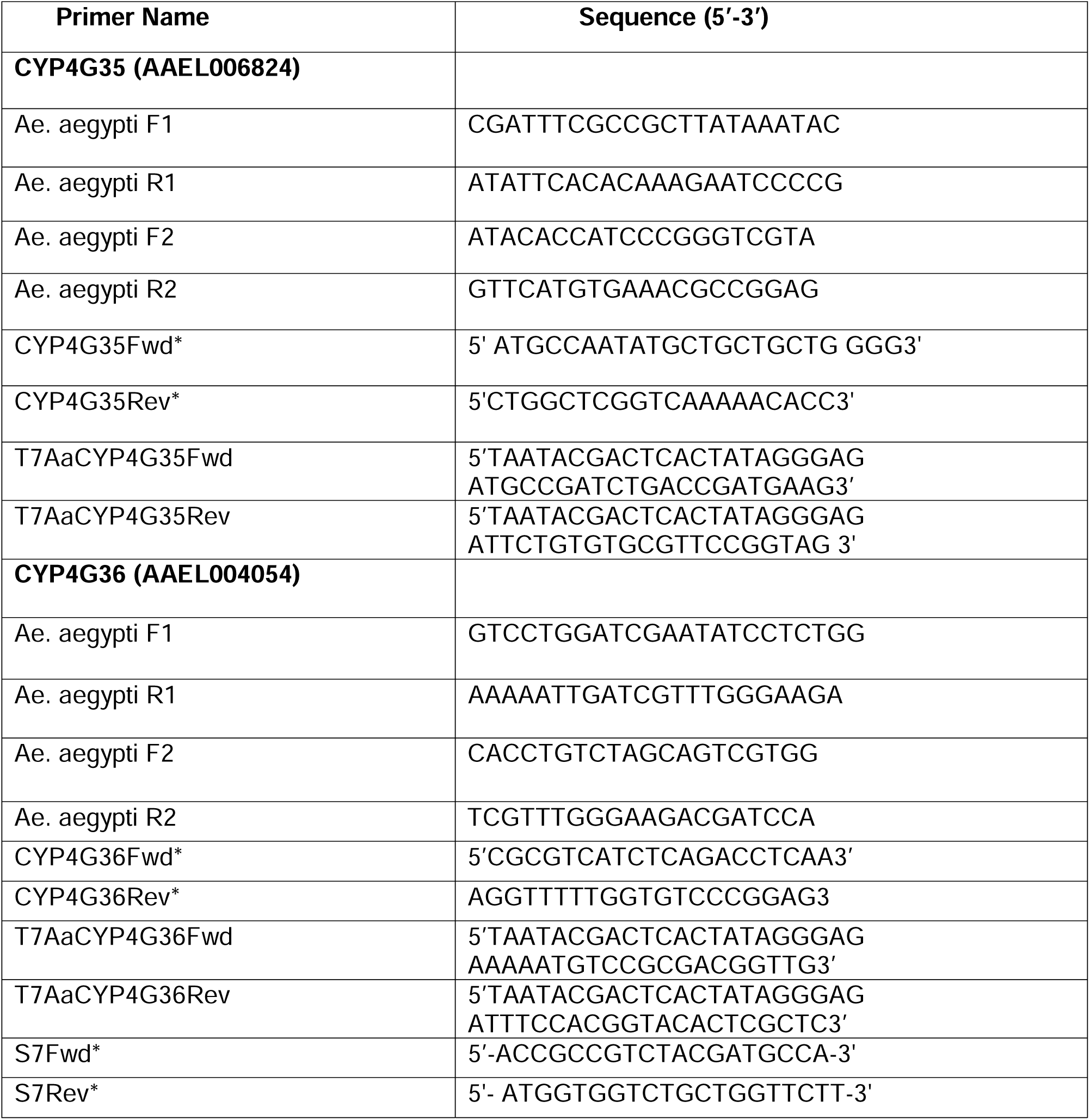
Primers for PCR amplify and sequence the CYP4G35 for transgenic lines. The internal primers set were primers: *Ae. aegypti* F2 and *Ae. aegypti* R2. The external primer set was primers: *Ae. aegypti* F1 and *Ae. aegypti* R1. * Primers used for knockdown validation of CYP4G35 and CYP4G36.

### dsRNA Preparation

Double-stranded RNA (dsRNA) templates were amplified by PCR using cDNA derived from head tissue. Gene-specific primers for CYP4G35 and CYP4G36 were designed with the T7 promoter sequence (Table 1). PCR conditions were 95 °C for 5 min, 95 °C for 30 s, 55 °C for 30 s, 72 °C for 30 s (5 cycles) followed by 95 °C for 30 s, 65 °C for 30 s, 72 °C for 30 s (35 cycles), and a final extension at 72 °C for 10 min. The PCR product was purified (Zymo PCR purification kit, Zymo Inc., Irvine, CA, USA), and 1 µg purified product was used for *in vitro* transcription. A non-specific gene, enhanced green fluorescent protein (EGFP) transcript was amplified from a plasmid. CYP4G35, CYP4G36, and EGFP dsRNA were prepared (Ambion Megascript T7 *in vitro* transcription kit, Austin, TX) as previously described^26,27^.

### RNA interference (RNAi)

Knockdown of CYP4G transcripts was performed as described in an earlier study^26^. Briefly, early pupae (5 pupae/well (n=3), within 4 h of molting) were placed in 12-well plates (3.9 cm^2^ growth area, Olympus Plastics, Genesee Scientific, El Cajon, CA, USA). Each well contained 5 µg (2 µl) of dsRNA (dsCYP4G35 or dsCYP4G36) per 500 µl of DNase-free water. Control wells contained 500 µl of water and EGFP dsRNA. Adult males and females were collected soon after eclosion (within 10 hours). Total RNA was extracted from CYP4G dsRNA-treated mosquitoes and the control, and cDNA was synthesized as described above. RT-PCR and gel electrophoresis were performed using CYP4G-specific primers to assess knockdown efficiency relative to control mosquito transcript levels. RT-PCR conditions were as follows: 95 °C for 2 min, 95 °C for 3 s, 52 °C for 30 s, 72 °C for 30 s (39 cycles), and a final extension at 72 °C for 10 min.

### Single guide RNA (sgRNA) design and construction

Three sgRNAs were designed for each CYP4G genomic DNA sequence by manually searching genomic regions for protospacer-adjacent motifs (PAMs) with the sequence “NGG”, where N is any nucleotide. Sequences of sgRNA were required to be 18–20 bp in length, excluding the PAM, and contained one or two 5′ terminal guanines to facilitate transcription by T7 RNA polymerase. sgRNA sequences were tested for potential off-target binding using the following two web tools: CRISPR (http://crispor.tefor.net/)^28^ and ChopChop (https://chopchop.rc.fas.harvard.edu/)^29^. Three sgRNA sequences were selected for each gene for the optimal efficacy (Table 2, Fig S1). sgRNAs were prepared using a PCR template amplified by CRISPR forward and reverse primers designed for each target gene and synthesized using short primer design and in vitro transcription (IVT) in the laboratory using the PCR template and GeneArt™ Precision gRNA Synthesis Kit (Thermo Fisher, Waltham, MA, USA) as per the manufacturer’s instructions.

**Table 2:**
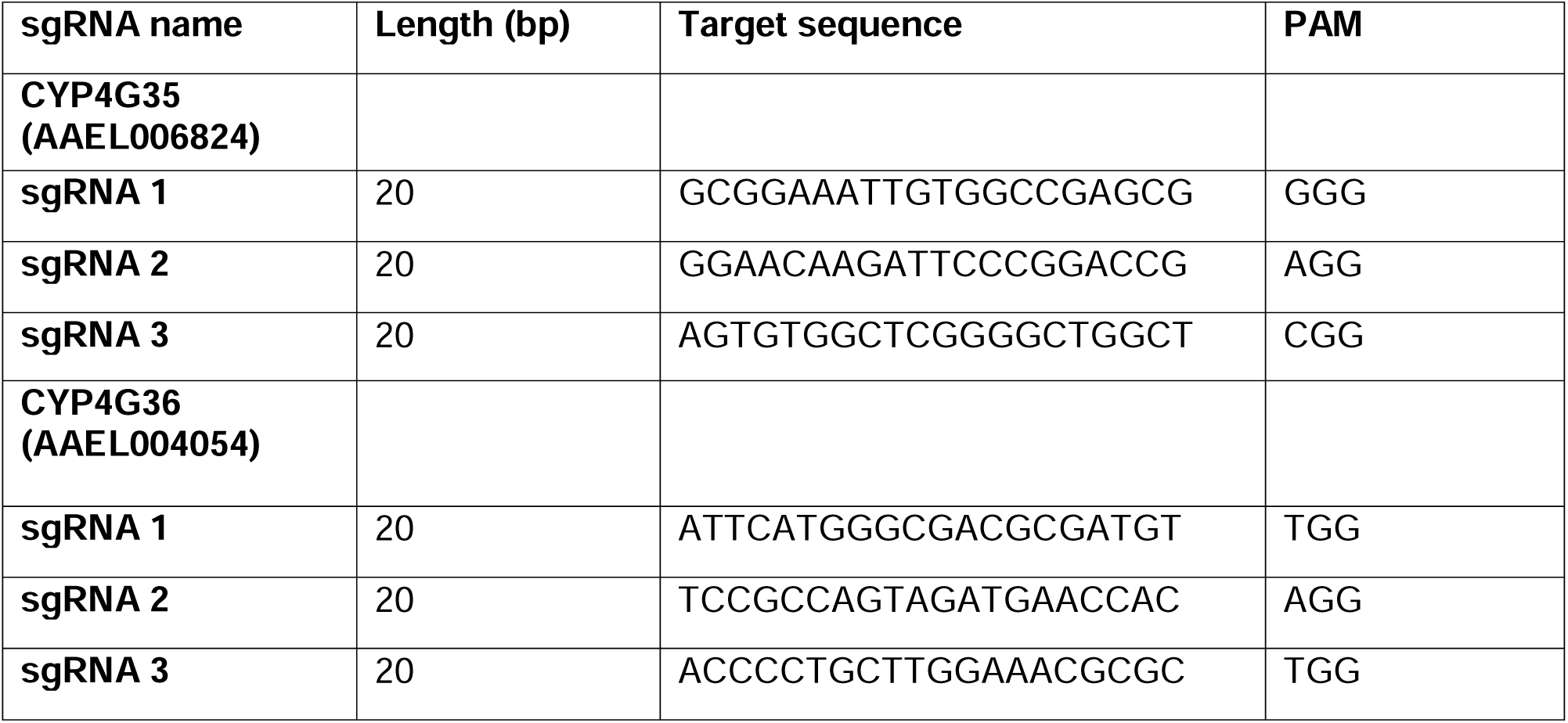
Guide RNAs designed used to target CYP4Gs in *Aedes aegypti*.

### Generation of CYP4G35 and CYP4G36 transgenic mosquitoes

The synthesized sgRNAs (a pool of three) were mixed (120 ng/μl) with injection-ready Cas9 protein (300 ng/μl: PNA Bio) in Cas9 Dilution Buffer (PNABio, Thousand Oaks, CA, USA)^30^. The sgRNA and Cas9 mix was sent to the Insect Transformation Facility at the University of Maryland and was injected into ∼250 pre-blastoderm-stage embryos. Injected embryos were allowed to recover for 3 days, then hatched under vacuum. The resulting G_0_ female adults were screened and backcrossed to wild-type (WT) males and individually set up to collect eggs for the next generation, which were screened again. G_0_ males were crossed with 5-10 WT females. Genomic DNA (gDNA) was extracted from half of a mosquito leg using Chelex^30,31^ as follows: mosquitoes were anesthetized in a 4 °C before leg removal. The tissue sample was placed in a PCR tube containing 200 μl of 10% Chelex 100 resin: water (v/v) (Sigma-Aldrich, CAS #: 11139-85-8), ensuring the sample was embedded in the Chelex portion of the solution. The PCR tubes containing the sample were then heated to 95 °C. The samples were vortexed for 5 s and centrifuged for 15 s. The supernatant was used for nested PCR to amplify regions surrounding the putative CRISPR-Cas9 cut site from the genomic DNA (gDNA) of a given generation, using primers listed in Table 3. All reactions were performed with an initial 2 min at 95 °C, followed by 39 cycles of 30 s at 95 °C, 30 s at 52 °C, and 30 s at 72 °C and a final extension at 72 °C for 10 min. PCR reactions were run on a 1.2% agarose gel to confirm a positive PCR amplification. PCR products were sent to UNR Genomics Center for PCR clean-up and Sanger sequencing using a set of internal primers to screen progeny for mutants. The sequences from injected individuals were aligned to the wildtype (WT) reference sequence and examined for insertions, deletions, or other polymorphisms. Synthego Inference of CRISPR Edits (ICE) analysis was employed to determine the contribution of gene editing and sgRNA ^32^.

**Table 3.**
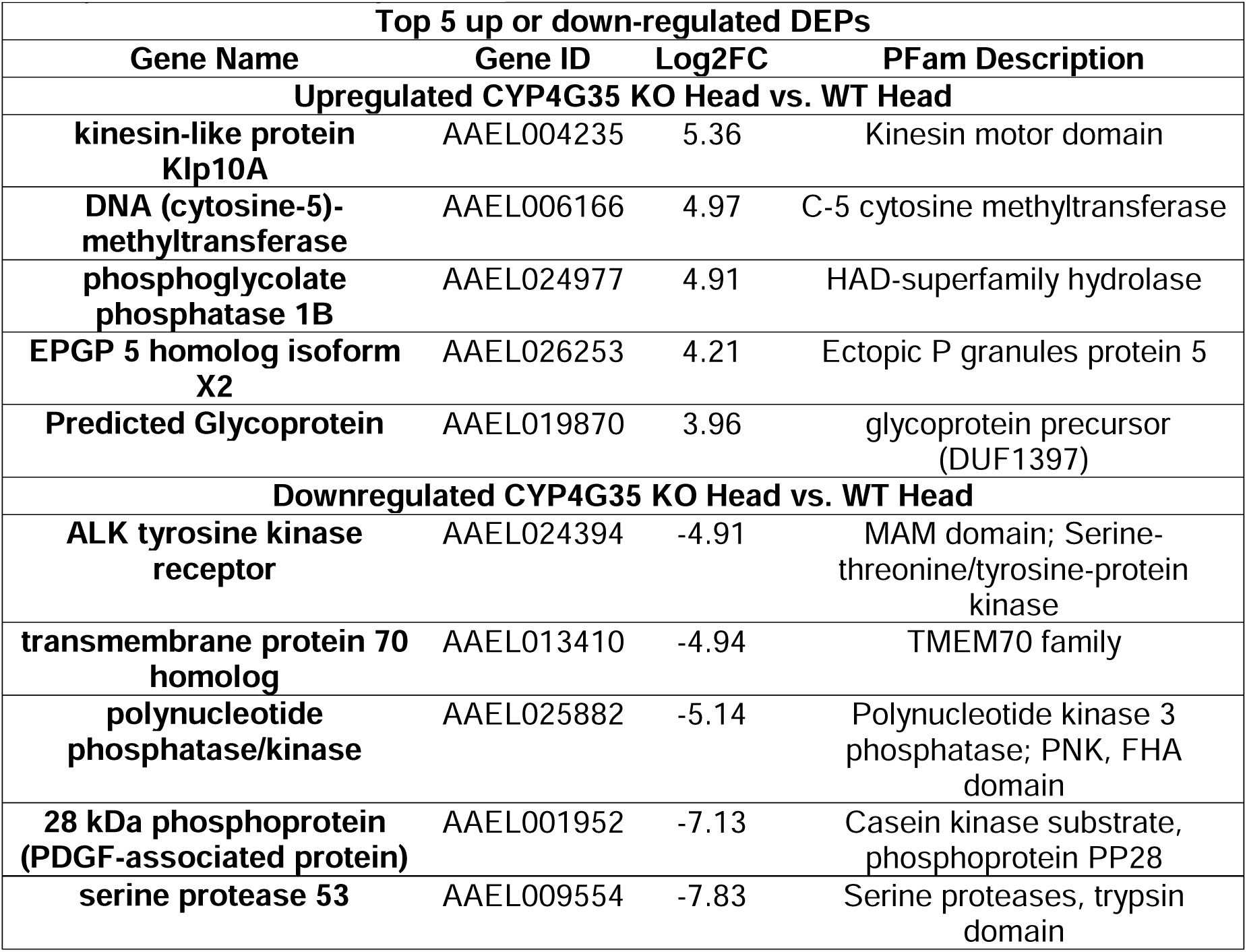
Top five up or downregulated differentially expressed proteins in CYP4G35 KO head vs. WT head. DEPs from Gene Name, Gene ID, Fold change (Log2FC), and protein family the transcript belongs to.

### Y-Tube olfactometer assay

The knockout mosquitoes were tested for their olfactory variation to human odor using a Y-tube olfactometer, which consisted of a cylindrical chamber with two arms measuring 26 cm, 21 cm, and 21 cm, respectively, and an inner diameter of 5 cm^33^. For each experiment, approximately 20-25, 4-5-day-old mated, non-blood-fed females were tested for their preference towards human odor. The female mosquitoes were placed in a release chamber and allowed to rest and acclimate for 20 min, before entering the central cylinder and making a choice. The females flew upwind against an airflow of 2.5 L/min and entered one of the two arms with a trapping chamber at the end. The trapping chamber (Y-arms of the olfactometer) was connected to a stimulus tube delivering either the control odor or the human odor. The experiment was concluded after 50 min (20 min acclimation and 30 min of choice).

The human odor was generated by wearing a white cotton stockinette on the volunteer’s leg for 10-12 hours. The control consisted of the stockinette without human odor. To account for potential spatial effects, all the tubing was cleaned with 70% ethanol and the location of the control and treatment was alternated every time the mosquitoes were introduced. The olfactometer was covered with a black cloth to prevent visual cues from influencing the experiment. Mosquitoes were allowed to find a “host” for 30 min, and those that reached the trapping chamber were considered to have made a choice, while those that remained in the acclimatization area or the long tube were considered to have no preference. The experiment was repeated three times with different cohorts of females, resulting in a total of approximately 75 mosquitoes per gene. The percent response was calculated as (total number of mosquitoes showing response) / (total number of mosquitoes used) *100.

### CYP4G Recombinant Fusion Proteins

CYP4G35 and CYP4G36 recombinant fusion proteins (with cytochrome P450 reductase, CPR) were expressed in Sf9 insect cells using the BaculoDirect™ expression system (Invitrogen), following the methods outlined by^19^ (Fig. S2). DNA inserts from the pENTR4 vector, which lacked the NcoI site, were transferred into the BaculoDirect™ C-terminal destination vector. Subsequently, P3 high-titer viral stocks were obtained through a series of amplification steps. Sf9 cells, seeded at a density of 1×10□ cells/mL, were infected with these recombinant viruses and cultured for 72 h in Sf900 II medium (Gibco). The medium was enriched with 10% fetal bovine serum (Atlas Biologicals), 0.3 mM δ-aminolevulinic acid, and 0.2 mM ferric citrate to support protein expression.

CYP4G55 from Pine beetle, *Dendroctonus ponderosae*^19^ was used as a positive control, and housefly CPR (HF-CPR) served as the negative control. Cells expressing CYP4G35-CPR, CYP4G36-CPR, CYP4G55-CPR, and HF-CPR were disrupted by sonication (4 watts, 1 second on/off, repeated 40 times) in a buffer containing 100 mM sodium phosphate (pH 7.6), 20% glycerol, 1.1 mM EDTA, 100 μM DTT, 200 μM PMSF, and protease inhibitors.

Following lysis, cell debris was removed by centrifugation at 10,000×g for 20 min at 4□°C. Subsequently, microsomal fractions were isolated through ultracentrifugation at 150,000×g for 90 min at 4□°C. Protein quantification was conducted using carbon monoxide difference spectroscopy^19^.

### Carbon monoxide (CO) difference spectrum

CYP4G-CPR-containing microsomes were assessed for functional activity as a measure of proper folding and a peak at 450 nm by CO-difference spectrum analysis. A spectrophotometer (VWR, SpectraMax® M5/M5e Multimode Plate Reader) measuring 400 to 500 nm was used to observe a Soret band absorption from the heme group after incubation with CO gas. Denatured CYP4G has only one Soret peak at 420 nm when saturated with CO. Fully reduced CYP4G has an additional Soret peak at 450 nm in its absorption spectra. In duplicate, microsomes were added to a 96-well plate (UV half-star plate). One of the two microsome samples was sealed to prevent CO gas from entering the well. The plate was then placed inside a sealed container, with the opening adjusted to connect to the CO gas tank. The CO valve was opened to release gas into the container and the wells for 2-3 min at 20-30 PSI. The plate was removed from the container, and the sealant protecting one of the wells was removed. To each well, 5 µL (0.5 M) of sodium dithionate was added and mixed. The plate was analyzed with a spectrophotometer at 5 min intervals (5, 10, and 15 min). Microsome preparations containing CYP4G35-CPR, CYP4G36-CPR, and CYP4G55-CPR were flash frozen and stored at -80 □C for up to 4 weeks.

### Substrate Specificity Assays

Each reaction contained 25 pmol of microsomal CYP, assay buffer (1.2 mM MgCl_2_, 1.2 mM CaCl_2_, 0.4 mM DTT), 25 μg substrate (n-heptadecane, decan-1-ol, and decanal), and 1.0 mM NADPH with a total reaction volume of 500 μl. The reactions were incubated overnight at 30 °C in a hybridization chamber and quenched with 200 μl of 2 M HCl. Approximately 20 mg of NaCl was added to each reaction, and products were extracted with 500 μl hexane: ether (1:1). Extracted products were analyzed using a Shimadzu QP2020 gas chromatograph-mass spectrometer with a molecular weight scanning range of 50 - 550 atomic mass units (AMU). A trace chromatograph containing a 30m, 25 μm film thickness DB-5 capillary column (Shimadzu Scientific Instruments, Columbia, MD, USA) was programmed with the following parameters: an initial temperature of 40 □C for 10-carbon-chain-long substrates with no hold, then programmed at 5 □C per minute to 25 □C.

### Antibody generation

Antibodies for both CYP4G35 (XP_001658068.2: antigen: KLPSPSLSEIIAKEESESKESLP-Cys (24aa) and CYP4G36 (XP_001648376.1: antigen: KTAEFEKPKSNINTNSVEGLS-Cys (22aa) were generated by Pacific Immunology (Pacific Immunology, Ramona, CA, USA; National Institutes of Health animal welfare assurance No. A41820-01). Synthetic peptides were used with complete Freund’s adjuvant (CFA) and incomplete Freund’s adjuvant (IFA) to immunize two rabbits per peptide. Pre-immune sera (5 ml.) were collected from all four rabbits. The first immunization with CFA was conducted on the same day as the collection of pre-immune sera. After that, two booster doses of IFA were given at three-week intervals each. Sera were collected for ELISA one week after the second booster, and again at 1 and 2 weeks later (Production bleed). A third and final booster was administered four weeks after the last booster, and rabbits were bled weekly for four weeks. Rabbits were euthanized according to IACUC protocol-21-01-118-2.

### Dot Blot Assay for detection of CYP4Gs antibodies

Dot blots were used to examine the specificity of antisera to mosquito CYP4Gs. For protein extraction, mosquitoes were collected in PBS, and a protease inhibitor (1X PBS-PI) solution, homogenized using a micropastle, and centrifuged at 12,000 x g for 5 min at 4 °C. The supernatant was collected and stored at 4 °C. 2 μL of protein was spotted onto a 0.2 μm nitrocellulose membrane (Thermo Fisher Scientific, Waltham, MA USA) and allowed to dry at room temperature (RT). The membrane was rewetted in Tris-buffered saline-1% Triton X-100 (TBS-T) (250mM Tris, 27mM KCl, 1.37M NaCl, pH 7.4) and blocked with blocking buffer (5% dry milk in TBS–T, 0.03% Triton X-100). Membranes were then incubated in primary antiserum diluted in 5% Bovine serum Albumin (BSA)-TBS-T (1/500, 1/1000, 1/5000, 1/25000) for 1 h at RT. The membrane was rinsed in TBS-T three times, 5 min each at RT, and incubated with the anti-rabbit conjugated secondary antibody, Alexa Fluor 647 (1:10,000 dilution) (Jackson ImmunoResearch Labs, West Grove, PA, USA), for 1 h at RT, followed by three washes in TBS-T for 5 min each. The membrane was scanned on an Odyssey Scanner (Li-Cor Biotechnology, Lincoln, NE, USA) at 700–800 nm for visualization.

### DIG-labeled RNA probe construction

Primers were designed to amplify a 584-nucleotide sequence, a region unique to CYP4G35. The resulting PCR product was cloned into the pMiniT 2.0 vector using NEB® PCR Cloning Kit. The pMiniT2.0 cloning vector contains T7 and SP6 polymerase priming sites, enabling the production of sense and antisense run-off transcription products from a single clone. The cloned amplicon was sequenced to verify the sequence fidelity and orientation within the vector plasmid. Restriction digestion was performed with BamHI and NotI to produce antisense and sense-strand RNA probes. Gel electrophoresis was used to verify that the pMiniT2.0 clone was linearized entirely. The linearized plasmid DNA was purified (Zymo PCR purification kit). The digoxigenin (DIG)-labeled single-stranded RNA probe was prepared by performing an *in vitro* transcription reaction using 1 µg of the linearized plasmid DNA as a template. The reaction was prepared with SP6/T7 RNA polymerase, transcription buffer, and DIG RNA labeling mix as described by the manufacturer’s instructions (Roche, DIG RNA labeling Kit (SP6/T7, SIGMA-Aldrich, Burlington, MA, USA)^34^. A total of 75 ng/ml of DIG-labelled mRNA was made for the antisense and sense probes.

### Whole-mount Immunohistochemistry (WM - IHC)

Heads of six-day-old sugar-fed females (8 heads; containing a pair of antennae) were removed from cold anesthetized mosquitoes using fine forceps and transferred directly to zinc formaldehyde (0.25% ZnCl2, 1% formaldehyde, 135 mM NaCl, 1.2 % sucrose, 0.03% Triton X-100) for 24 h^35^. Following fixation for 24 h at RT, antennae were washed with Hepes Buffer Saline (HBS) for 15 mins (150 mM NaCl, 5 mM KCl, 25 mM sucrose, 10 mM Hepes, 5 mM CaCl2, 0.03% Triton X-100). The heads were transferred into a concave glass slide containing HBS buffer and observed under a microscope using ultra-fine forceps. The antennae were removed from the head, carefully squeezed ten times with ultra-fine forceps, and then transferred to PCR tubes. The antennae were incubated in an HBS buffer for 15 min and in 80% methanol / 20% DMSO^35^. Subsequently, the antennae were incubated in a 5% BSA blocking solution (PBS, 5% BSA, 1% DMSO, 0.03% Triton X-100) overnight. The solution was replaced by a 5% BSA blocking solution containing CYP4G35 or CYP4G36 primary antibodies (dilution 1:100), and the PCR tube was placed for 30 s in a water bath sonifier (FS30 Ultrasonic Cleaner, Fischer Scientific) at 40 Hz. Antennae were incubated for four days at 6 °C, with sonication repeated on the second day. This was followed by washing thrice for 15 min each at RT in PBS, 1% DMSO, and 0.03% Triton X-100. The secondary antibody (anti-Rabbit Alexa Fluor 555) was diluted (1:1000) in blocking solution and added to the samples. After incubation, the samples were washed three times for 15 min each in PBS with 1% DMSO, 0.03% Triton X-100, and a short rinse in PBS. Once placed on charged slides, the antennae were mounted in Mowiol (Sigma-Aldrich, Burlington, MA, USA) mounting media. The coverslip was then placed over the sample for 24-48 h before sealing it with nail polish. All experiments were repeated thrice with a different cohort of mosquitoes.

### Whole-mount Fluorescent *in-situ* hybridization (WM-FISH)

Female *Ae. aegypti* heads were dissected from cold anesthetized six-day-old, sugar-fed mosquitoes and transferred directly to zinc formaldehyde (as explained in WM-IHC protocol) for 24 h at RT for fixation^35^. Tissues (antennae, palps, and proboscis) were washed with HBS buffer and transferred onto a concave glass slide. Under a microscope using ultra-fine forceps, the samples were squeezed, with forceps holding one end of the tissue, and carefully washed in HBS buffer for 15 min and incubated in 80% methanol, 20% DMSO (as explained in WM-IHC protocol)^35^. The tissues were transferred to a PCR tube and then incubated in prehybridization solution (50% formamide, 5x SSC, 1x Denhardt’s reagent, 50 μg/ml yeast RNA, 1% Tween 20, 0.1% Chaps, 5 mM EDTA, pH 8.0) at 45 °C for 5–6 h and followed by incubation for three days at 55 °C in the same solution containing a DIG-labeled antisense RNA at (1:10-1:100). In control experiments, DIG-labeled sense RNA probes instead of antisense RNA probes were used. Once prehybridization was completed, the tissues were washed in 0.1 x SSC, 0.03% Triton X-100 four times for 15 min each, and treated with a blocking solution^35^ followed by the blocking solution containing the anti-DIG Alexa Fluor 647 (1:1000) (Jackson ImmunoResearch Labs, West Grove, PA, USA,). Subsequently, the tubes were placed for 30 s in a water bath sonifier (FS30 Ultrasonic Cleaner, Fischer Scientific) and incubated for 3 days at 6 °C, washed three times for 15 min each in PBS with 1% DMSO, 0.03% Triton X-100. Samples were transferred to charged slides, mounted in Mowiol mounting medium, and sealed with nail polish. All experiments were repeated thrice with a different cohort of mosquitoes.

### Microscopy

Tissue samples from WM-IHC and WM-FISH experiments were analyzed for fluorescence using a BZ-X800 Keyence microscope (Keyence, United States). Images of the red (protein) and green (mRNA probe) fluorescence channels were recorded. All images were captured at 10x, 40x, and 100x magnification.

### Proteomics sample collection and data analysis

For proteome samples, WT and CYP4G35-KO adults were collected from six-day-old, sugar-fed females. Head and body were separated from 20 WT (WT-Head, WT-Body) and CYP4G35KO (KO-Head, KO-Body) mosquitoes each (N=3). Tissues were washed in cold 1X PBS (Santa Cruz Biotechnology, Dallas, TX, USA) containing one PierceTM Protease Inhibitor tablet (Thermo Fisher, Rockford, IL, USA) (PBS+PI). Tissue samples were transferred to 1.7 ml microtubes containing 200 µl PBS+PI and homogenized using microtube pestles. After homogenization, 400 µl of PBS with PI was added to the samples, followed by centrifugation at 12,000 × g for 10 min. The supernatant was then transferred to a new microtube and centrifuged at 12000 x g. Finally, the supernatant was collected and sent to the UNR Proteomics Core (University of Nevada, Reno).

#### Protein digestion

The protein content of the prepared samples was estimated using the fluorescence-based protein assay EZQ (Invitrogen #R33200). During enzymatic sample processing, 50 μg of protein extracts were reduced, alkylated with iodoacetamide, and digested according to the protocol provided in the Thermo Scientific EasyPep Mini MS Sample Prep Kit (Cat #A40006). The trypsin/Lys-C was added at a 1:10 (enzyme: protein) ratio for digestion. At this point, the supernatant consisted of peptides and contaminants. The supernatant was placed in the Peptide Clean-up column provided, which was used to remove impurities through various washes using the Mini MS Sample prep kit (Cat #A40006). Following the instructions provided in the kit, the multiple flow-throughs would remove both hydrophilic and hydrophobic contaminants. The sample was eluted from the column supplied with the kit. The peptides were dried in a vacuum centrifuge and resuspended in 100 μl of 0.1% formic acid for LC-MS analysis.

#### Liquid Chromatography

Samples were analyzed using an UltiMate 3000 RSLCnano system (Thermo Scientific, San Jose, CA). The peptides were trapped before separation on a 300 µm i.d. × 5 mm C18 PepMap 100 trap (Thermo Scientific, San Jose, CA) for 5 min at 10 μL/min. Next, separation was performed on a 50 cm uPAC C18 nano-LC column (PharmaFluidics, Ghent, Belgium) using an EasySpray source (Thermo Scientific, San Jose, CA) equipped with a 30-μm ID stainless steel emitter (PepSep, Marslev, Denmark). Separation was performed at 350 nl/min using a gradient from 1% to 45% for 60 min (Solvent A: 0.1% Formic Acid, Solvent B: Acetonitrile, 0.1% Formic Acid).

#### Mass spectrometry

Data-Independent Acquisition (DIA)^36,37^ was performed using a chromatographic library ^38,39^, using an Eclipse Tribrid Orbitrap mass spectrometer (Thermo Scientific, San Jose, CA). Briefly, six gas-phase fractions (GPFs) of the biological sample pool were used to generate a reference library. The GPF acquisition used 4 m/z precursor isolation windows in a staggered pattern (GPF1 398.4-502.5 m/z, GPF2 498.5-602.5 m/z, GPF3 598.5-702.6 m/z, GPF4 698.6-802.6 m/z, GPF5 798.6-902.7 m/z, GPF6 898.7-1002.7 m/z) at a resolution of 60,000, AGC target 4e5, maximum injection time 55 ms, and an NCE of 33 using higher-energy collision dissociation (HCD). Biological samples were run on an identical gradient as the GPFs using a staggered window scheme of 24 m/z over a mass range of 385-1015 m/z. Precursor isolation was performed at 60,000 resolution, maximum injection time of 55 ms, AGC target 4e6, and an NCE of 33 using HCD. An empirically corrected library, which combines the GPF and the deep neural network Prosit^38^, was used to generate predicted fragments and retention times.

ScaffoldDIA v-3.1.0^40^ was used to process the experimental samples, allowing only peptides that were exclusively assigned to a given protein to be used for quantitative analysis and requiring a minimum of two peptides with a protein-level FDR of less than 1%. DIA-MS data files were converted to mzML format using ProteoWizard (3.0.19254). Deconvolution of staggered windows was performed. The intermediate chromatogram library was created by Encyclopedia (1.2.2). Reference samples were individually searched against (VectorBase62_Aaegypti LVP_AGWG_ AnnotatedProteins), containing 28,391 sequences with a peptide mass tolerance of 10.0 ppm and a fragment mass tolerance of 10.0 ppm. Fixed modifications considered were: Carbamidomethylation C. The digestion enzyme was assumed to be Trypsin, with a maximum of 1 missed cleavage site(s) allowed. Only peptides with charges in the range [2…3] and lengths in the range [6…30] were considered. Peptides identified in each search were filtered by Percolator (3.01. nightly-13-655e4c7-dirty) to achieve a maximum false discovery rate (FDR) of 0.01. Individual search results were combined, and peptides were again filtered to an FDR threshold of 0.01 for inclusion in the reference library. Individual search results were combined, and peptide identifications were assigned posterior error probabilities and filtered to an FDR threshold of 0.01 by Percolator. Peptide quantification was performed using EncyclopeDIAia^39^, a library search engine that enables peptide identification using DIA-based chromatogram libraries. For each peptide, the 5 highest-quality fragment ions were selected for quantitation. Proteins that contained similar peptides and could not be differentiated based on MS/MS analysis were grouped to satisfy the principles of parsimony. Protein groups were a threshold to achieve a protein FDR of less than 1.0%.

### Gene Ontology (GO) and pathway enrichment analysis of differentially expressed proteins (DEPs)

GO annotations were performed using the DAVID database^41^. GO terms were analyzed, and the primary process of GO analysis included BLAST, GO mapping, GO annotation, and InterProScan for annotation augmentation. Protein differences were calculated using Fisher’s exact test. GO enrichment of differentially expressed proteins was analyzed for biological processes, cellular components, and molecular functions using Blast2Go (Software Tool, https://www.blast2go.com).

KEGG pathway enrichment analysis was performed using the online analysis database on VectorBase^42^. Up- and down-regulated DEPs were separately subjected to GO and pathway enrichment analysis. P value <0.05 was used as the threshold value. DEPs were clustered using the hierarchical cluster method, and the results were presented as heat maps.

### Protein-protein interaction (PPI) network analysis

The STRING database (http://string-db.org/) was used to identify the interactions between proteins encoded by DEPs based on experimental data ^43^. A combined score of >0.4 was set as the threshold value. PPI networks were constructed with Cytoscape software^44^.

## RESULTS

### *Ae. aegypti* genome has two CYP4Gs paralogs

A key characteristic of CYP4Gs is an extended loop between the G and H helices. On this basis, two CYP4G paralogs were identified in the *Ae. aegypti* genome: CYP4G35 (Accession No. XM_001658018.2) and CYP4G36 (Accession No. XM_001648326.2). The naming of these CYPs has been changed to CYP4G15; however, this name is already assigned to *D. melanogaster* ^12^, which causes confusion. To maintain clarity, we continue to use the original nomenclature: CYP4G35 (ortholog of *D. melanogaster* CYP4G15) and CYP4G36 (ortholog of *D. melanogaster* CYP4G1).

### Both CYP4Gs have the highest expression in pharate adults. CYP4G35 expression is highest in female antennae, whereas CYP4G36 is expressed ubiquitously

Expression of both CYP4G35 and CYP4G36 was detected in all life stages. However, expression levels were significantly lower during the aquatic larval stages than during the pharate and terrestrial adult stages (Fig. 1A, B). Notably, the expression of both CYP4G paralogs remains elevated in newly eclosed adults but reaches levels similar to those in the aquatic stages once the females have blood-fed (Fig. 1A, B). We did not check the expression in older males.

**Figure 1:**
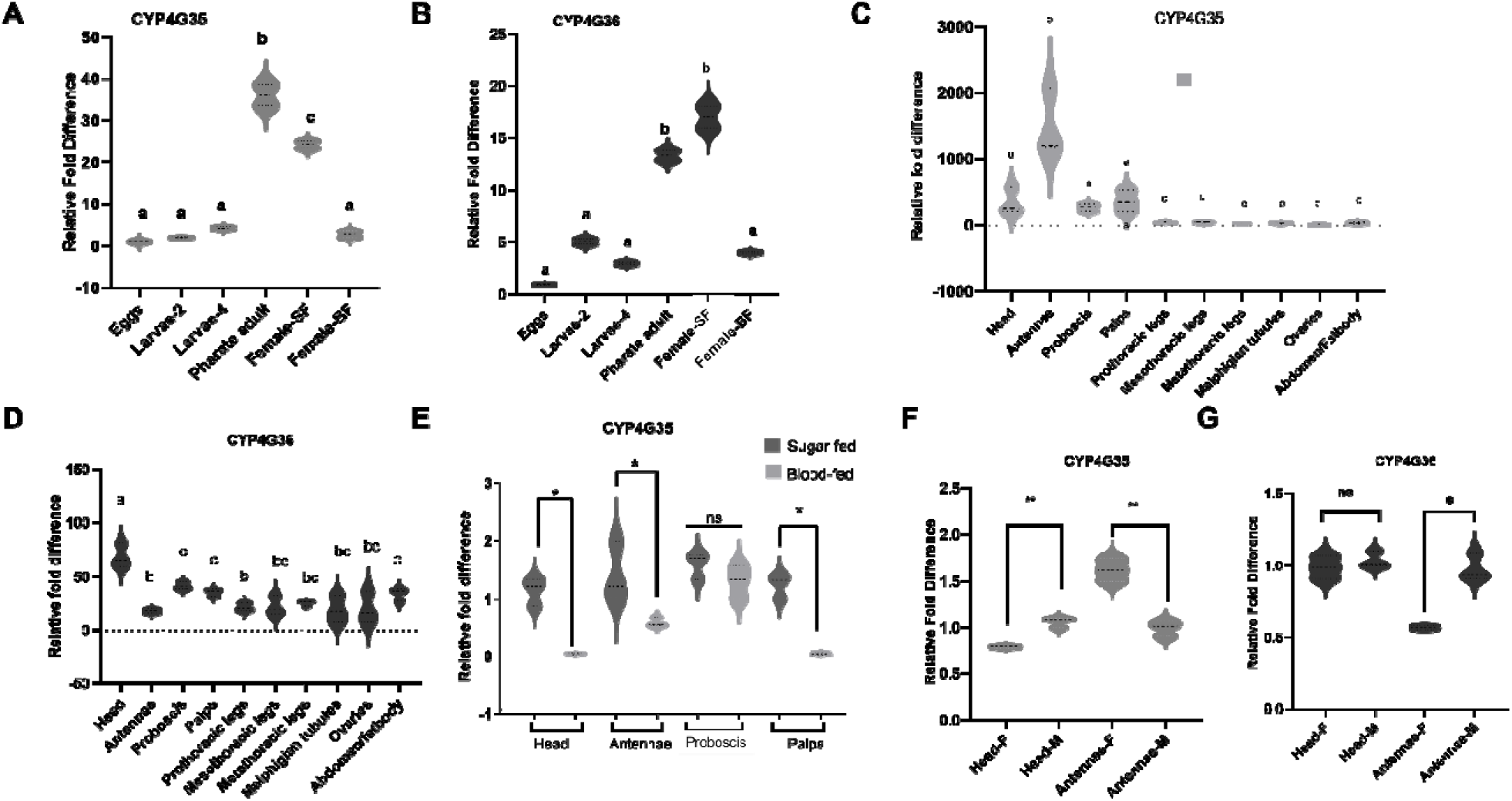
Temporal and Spatial expression of CYP4Gs. Transcript expression of CYP4G35 (A) and CYP4G36 (B) in developmental stages. qRT-PCR was performed to determine CYP4G expression levels across *Aedes aegypti* developmental stages. Egg samples were used as a control for relative expression for both CYP4G35 and CYP4G36. The data are means ± standard error. n=3. Transcript expression of CYP4G35 (C) and CYP4G36 (D) in female tissues. Expression levels of CYP4Gs in the Head, Antennae, Proboscis, Palps, pro, meso, and metathoracic legs, Malpighian tubules, Ovaries, and Abdominal wall/fat body were measured. Ovary samples were used as a control for relative expression. The data are means ± standard error. n=3. Difference in transcript expression of CYP4G35 in blood-fed and sugar-fed females (E). Sugar-fed females were used as a control for relative expression. n=3. Difference in transcript expression of CYP4G35 (F) and CYP4G36 (G) in male and female olfactory tissues. Head tissue contains mouth parts. Male tissue samples were used as a control for relative expression. n=3. One-way ANOVA and Kruskal-Wallis multiple comparison test were used for A-D. Welch’s t-test was used to compare two sets of data for E-G. SF- sugar-fed, BF-blood-fed, Hd- head, Ant- Antennae, F- female, M-male

In adult female mosquitoes, CYP4G35 expression was predominantly restricted to olfactory tissues, with the highest expression in antennae, followed by the head (containing mouthparts), proboscis, and maxillary palps. No expression was detected in non-olfactory tissues (Fig. 1C). In contrast, the highest expression of CYP4G36 was observed in the head; however, overall, this CYP exhibited a broader expression profile and was detected in all examined tissues (Fig. 1D).

Expression levels of CYP4G35 were significantly higher in sugar-fed females compared to their blood-fed counterparts in all tissues except the proboscis (Fig. 1E). CYP4G35 had higher expression in female antennae compared to males (Fig. 1F). In contrast, CYP4G36 showed significantly lower expression in female antennae (Fig. 1G), supporting the previous data in female tissue where CYP4G36 expression is lower in antennae (Fig. 1D).

### CYP4G35, not CYP4G36, localizes to the base and shaft of antennal sensilla

To further understand the localization of CYP4Gs in sensory tissue, antibodies against CYP4G35 and CYP4G36 were generated for immunohistochemistry (IHC), and mRNA probes were designed for fluorescent *in situ* hybridization (FISH). Antibody binding to antigens was confirmed by dot blots (Fig. S3). IHC with CYP4G35 antibody showed fluorescence in female antennal sensilla (Fig. 2A–B), absent in samples stained with CYP4G36 antibody or in secondary antibody-only controls (Fig. 2C-D). High-magnification images demonstrated its expression in antennal sensilla (base and shaft) (Fig. 2G), which, based on the shape, appear to be trichoid sensilla. While we fixed and imaged the entire antenna, the focus was on the first four antenna segments, which have previously been shown to be involved in host seeking ^45^. Fluorescence observed at antennal joints in all samples reflects autofluorescence or non-specific binding. Whole-mount FISH analysis using a DIG-labeled antisense probe for CYP4G35 also demonstrated mRNA localization at the base of the sensilla and along the shaft (Fig. 3A–B). Notably, no or very weak signal was observed in the sense probe and no-probe control samples (Fig. 3C–D). *In situ* antibody specificity has not yet been confirmed by KO tissue; these controls will be included in follow-up experiments.

**Figure 2.**
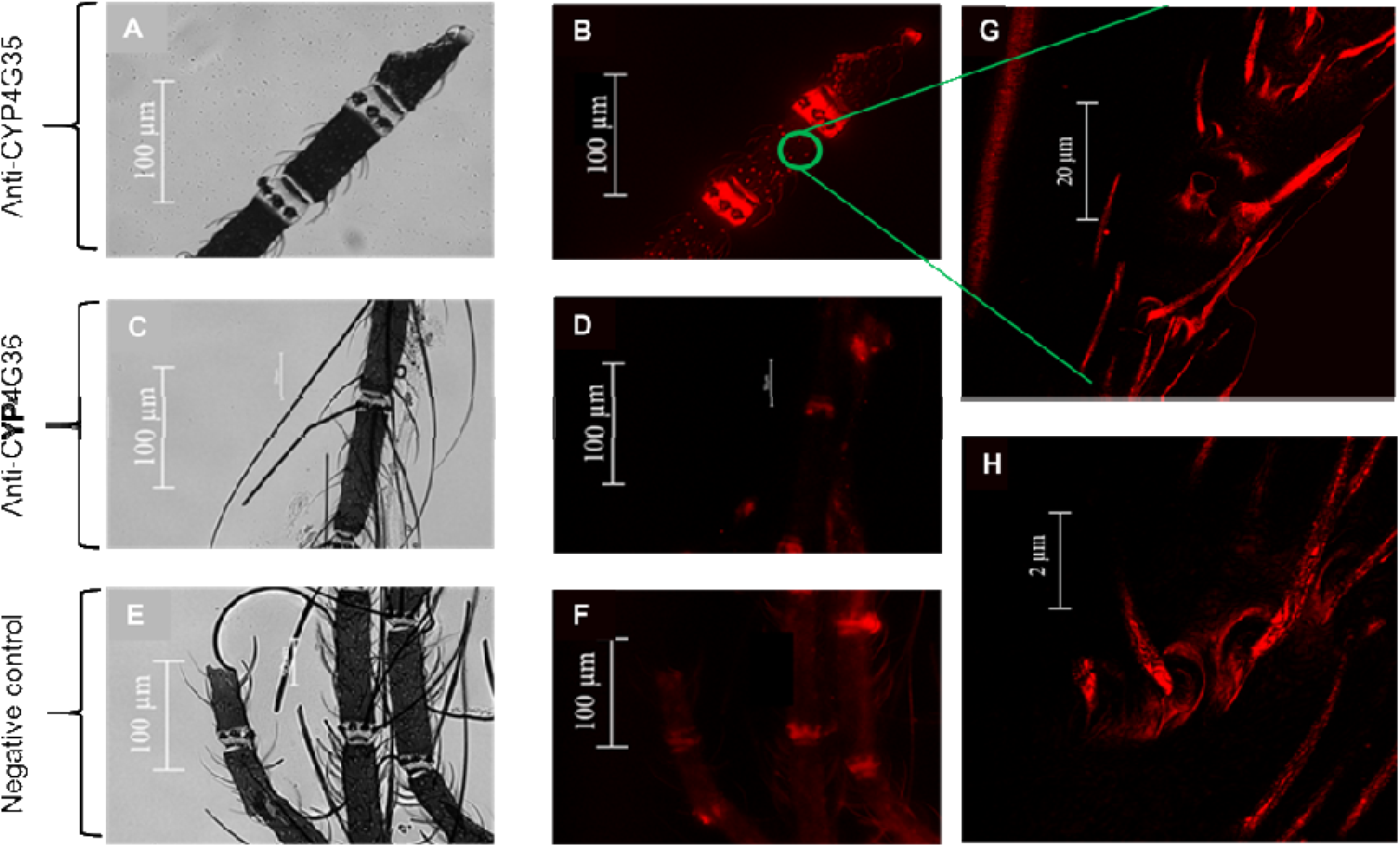
Localization of CYP4G35 and CYP4G36 in *Ae. aegypti* female antennae using Whole-mount fluorescent immunohistochemistry. Antennae were excised from the head using superfine forceps and fixed. Whole-mount fluorescent immunohistochemistry (WM-FIHC) was carried out using anti-CYP4G35 or anti-CYP4G36 primary antibodies and an anti-rabbit Alexa Fluor 555 secondary antibody. Representative bright field image of a CYP4G35 antibody-stained antenna (A), CYP4G35 antibody fluorescent staining (BZ-X Filter TRITC) (B). Bright field image of a CYP4G36 antibody-stained antenna tissue (D) and CYP4G36 antibody fluorescent staining (BZ-X Filter TRITC) (E). Only the secondary antibody AF 555 was used as control (G-H). Bright-field image of the antennae tissue (G) and secondary antibody AF 555 fluorescent staining (BZ-X Filter TRITC) (H). Images were taken on a Keyence BZ-X at 40x magnification. Magnified images of the sample in B were obtained using the SP8 confocal microscope (Lighting setting, Max resolution to speed setting) at 100x (I) and 125x (J).

**Figure 3.**
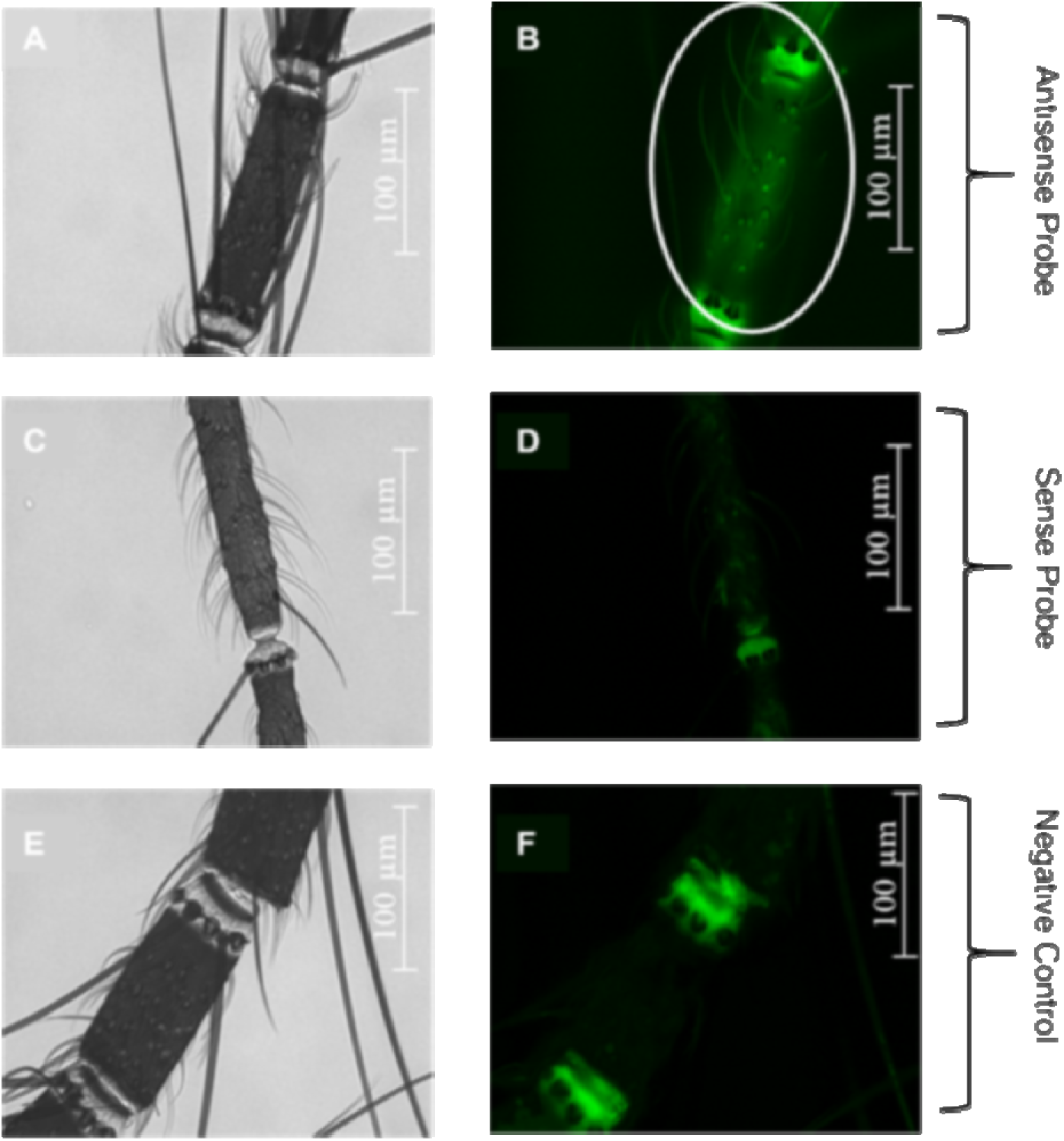
Localization of the CY4G35 in *Ae. aegypti* female antennae using whole-mount fluorescent *in-situ* hybridization. Antennae were excised from the head using superfine forceps and fixed. Whole-mount fluorescent *in situ* hybridization (WM-FISH) was carried out using digoxigenin-labeled (DIG) probes designed from a unique sequence of CYP4G35, and an anti-DIG Alexa Fluor 647A was used to detect the DIG signal. A sense probe was used as a control, and samples with only anti-DIG antibody were used as a negative control. A representative bright field image of the antenna tissue (A) and CYP4G35 antisense DIG-labeled probe fluorescent staining (BZ-X Filter CY5) (B). Bright field image of the antenna tissue labeled with CYP4G35 sense probe (C) and fluorescent staining (BZ-X) (D). Bright field image of the anti-DIG antibody only control (E) and fluorescent image (F). Images were captured on a Keyence BZ-X at 40x magnification.

In the proboscis, dual IHC-ISH demonstrated robust CYP4G35 expression at the sensilla-rich tip (Fig. 4A–D), which was absent in negative controls (Fig. 4E–H). In contrast, the maxillary palps did not exhibit any specific staining for either CYP4G35 or CYP4G36 (Data not shown).

**Figure 4.**
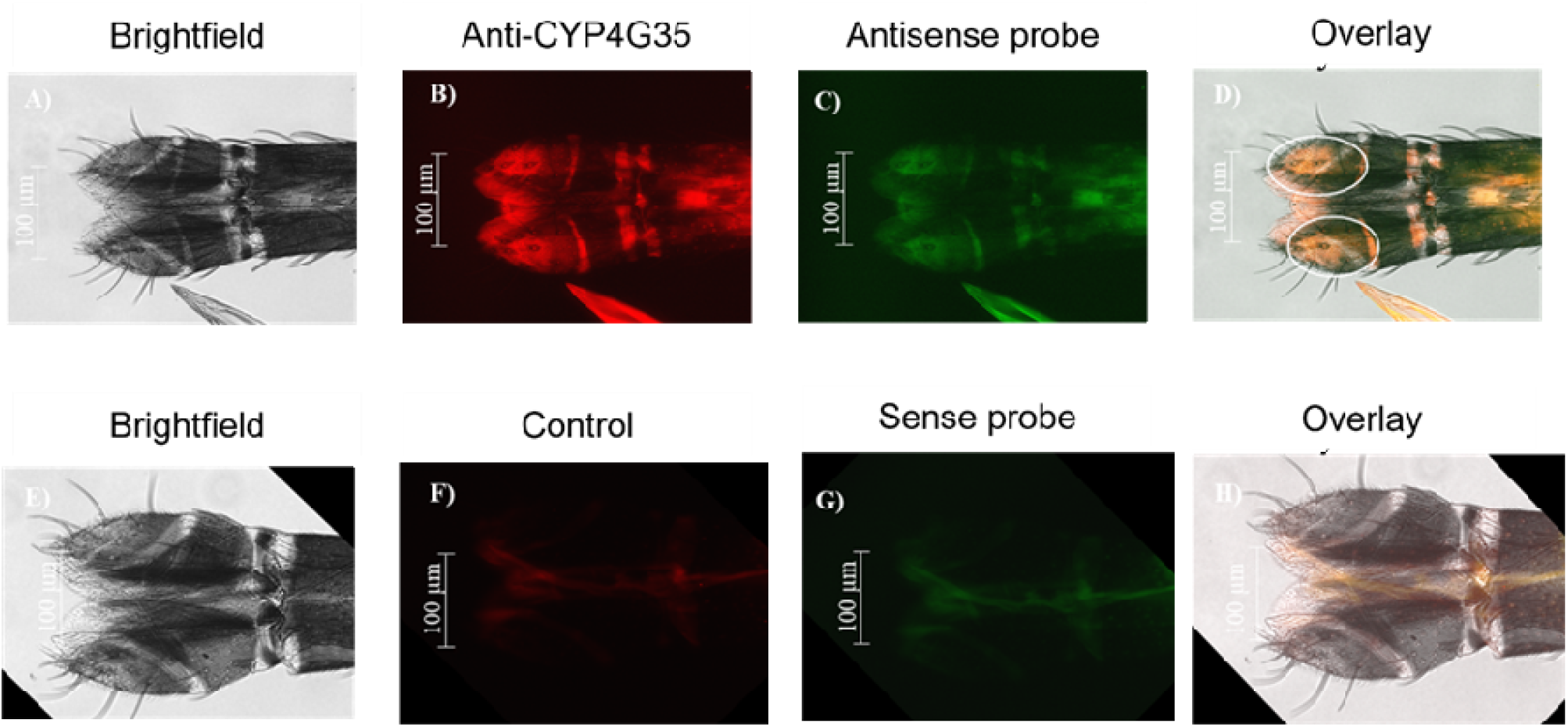
Localization of CYP4G35 in *Ae. aegypti* female proboscis using Whole-mount fluorescent *in-situ* hybridization and fluorescent immunohistochemistry. Proboscis were excised from the head using superfine forceps and fixed. Whole-mount fluorescent immunohistochemistry (WM-FIHC) was carried out using anti-CYP4G35 or anti-CYP4G36 primary antibodies and an anti-rabbit Alexa Fluor 555 secondary antibody. Whole-mount fluorescent *in-situ* hybridization (WM-FISH) used the CYP4G35 antisense DIG-labeled probe and an anti-DIG Alexa Fluor 647A was used to detect the DIG signal. All images were captured at 40x magnification (A-D). A) A representative brightfield image of the proboscis. B) CYP4G35 antibody staining (BZ-X Filter TRITC). C) CYP4G35 antisense DIG-labeled probe fluorescent staining (BZ-X CY5 filter). D) Overlay image of images A-C. E) Brightfield image F) No primary CYP4G35 antibody, secondary antibody only control, G) CYP4G35 sense DIG-labeled probe fluorescent staining (BZ-X CY5 filter). H) Overlay image of images E-G. The BZ-X Keyence microscope captured images. Post-capture image analysis was performed using BZ-X analysis software.

### CYP4G36 displays oxidative decarbonylase activity in our microsomal assay; CYP4G35 showed no detectable activity with the tested substrates

Most CYP4G enzymes characterized to date catalyze the oxidative decarbonylation of long-chain fatty alcohols, such as heptadecanol, to form aldehydes or hydrocarbons, supporting their role in cuticular hydrocarbon synthesis ^14,19,46^. To investigate the functions of *Ae. aegypti* CYP4Gs, recombinant CYP4G35 and CYP4G36 were expressed as fusion proteins with the catalytic domain of housefly cytochrome P450 reductase (HF-CPR). Mountain pine beetle CYP4G55 and honeybee CYP4G11 were included as controls, as they are known to utilize both short- and long-chain substrates ^13,19^. HF-CPR was used as a negative control. A functional protein would bind to CO and be detected at a 450 nm peak, corresponding to the heme group in the microsome. All fusion proteins exhibited characteristic CO-difference spectra with absorbance at 450 nm (Fig. 5A, B, C), indicating proper folding and NADPH reduction, which suggests the formation of a functional fusion protein (Fig. 5D).

**Figure 5.**
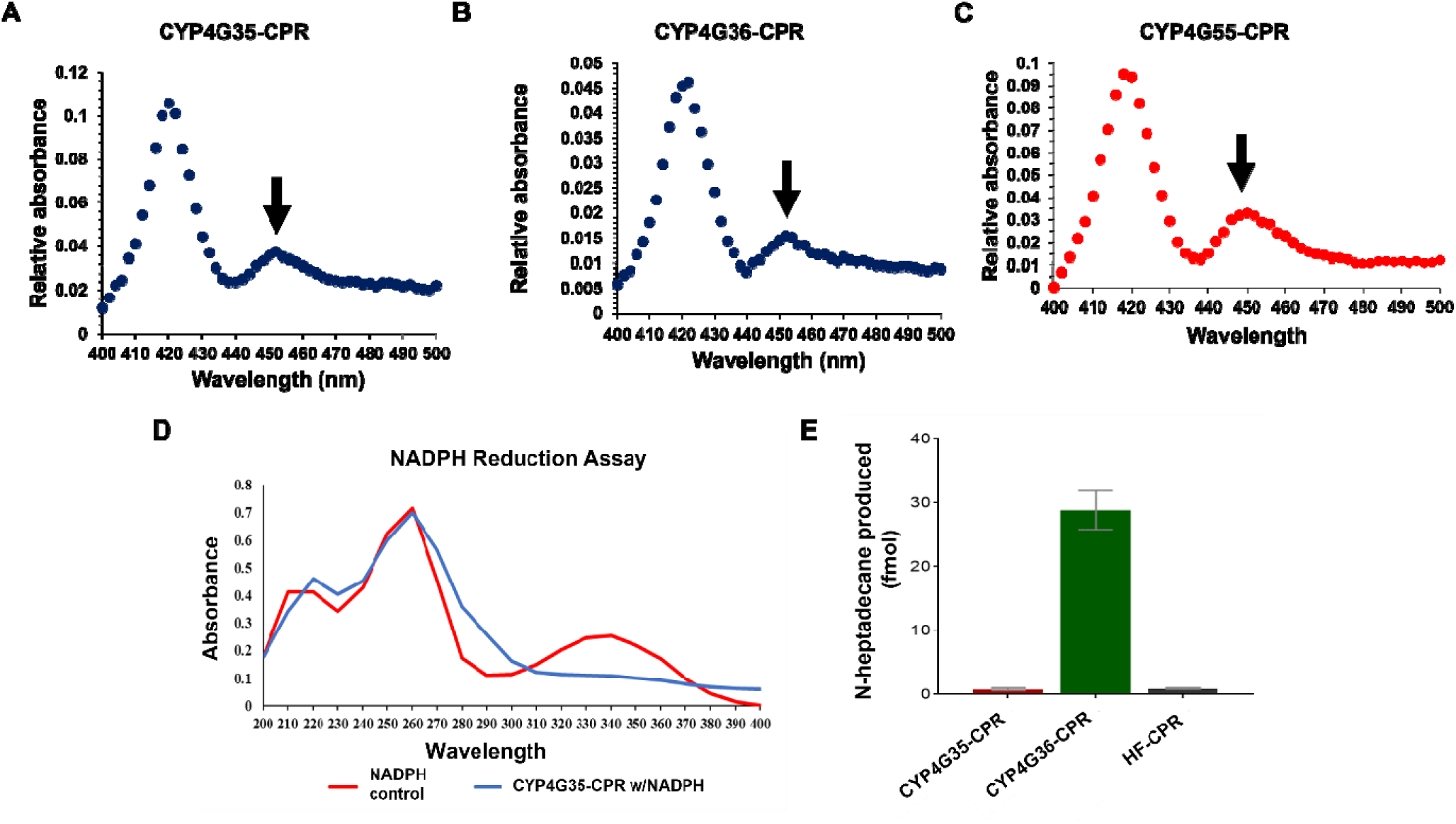
Validation of functional Cytochrome P450. Carbon monoxide (CO)-difference spectra of microsomes prepared from *Spodoptera frugiperda* (Sf)9 cells producing recombinant CYP4G35-CPR (A), CYP4G36-CPR (B), and *Dendroctonus ponderosae* CYP4G, CYP4G55-CPR (C). The fusion proteins produced the 450-nm peak (arrow) characteristic of functional cytochromes P450. CYP4G35-CPR activity was measured using NADPH reduction to NADP+, indicated by a decrease in absorbance at 340 nm. Compared to the NADPH control (D). The efficiency of CYP4Gs as an oxidative decarbonylase is determined by the conversion of octadecanal substrate to n-heptadecane product (E). CYP4Gs were produced as fusion proteins with house fly (HF) CPR. CPR- cytochrome P450 reductase.

CYP4G36-CPR efficiently converted [9,10-³H_₂_]-octadecanol to [8,9-³H_₂_]-n-heptadecane, yielding approximately 30 fmol of product per reaction. In contrast, CYP4G35-CPR and the CPR-only control (HF-CPR) showed minimal activity, producing less than 1 fmol per reaction (Fig. 5E). Attempts to test short-chain substrates failed to yield products.

### RNAi-mediated reduction of CYP4G35 is associated with delayed host-seeking but does not affect desiccation tolerance

Because CYP4G35 expression was highest in antennae and other olfactory tissues, we hypothesized that it may play a role in host seeking. We predicted that CYP4G35 knockdown mosquitoes would be impaired in detecting and responding to host odors, whereas CYP4G36 knockdown would have minimal or no effect on host seeking. To test this, we used RNAi. Pupal soaking resulted in approximately 80% knockdown of CYP4G35 and 65% knockdown of CYP4G36 in newly emerged adult females (Fig. 6A). In behavioral assays, CYP4G35-RNAi females failed to blood feed (Fig. 6B). Careful observation showed they did not orient themselves towards a blood source (human arm). In contrast, CYP4G36-RNAi females displayed only a slight delay in host-seeking in comparison to dsEGFP controls, with 100% feeding successfully within 20 min (Fig. 6B). Only 2 out of 20 CYP4G35 RNAi females probed after 15 min, and none fed to engorgement. In contrast, control and CYP4G36 RNAi females started probing within 1 min and by 15 min, all females were engorged. The CYP4G35 RNAi females (5/20) did feed when the arm was inside the cage and in direct contact with the mosquitoes; however, it took them ∼40 min to orient and feed (personal observation).

**Figure 6:**
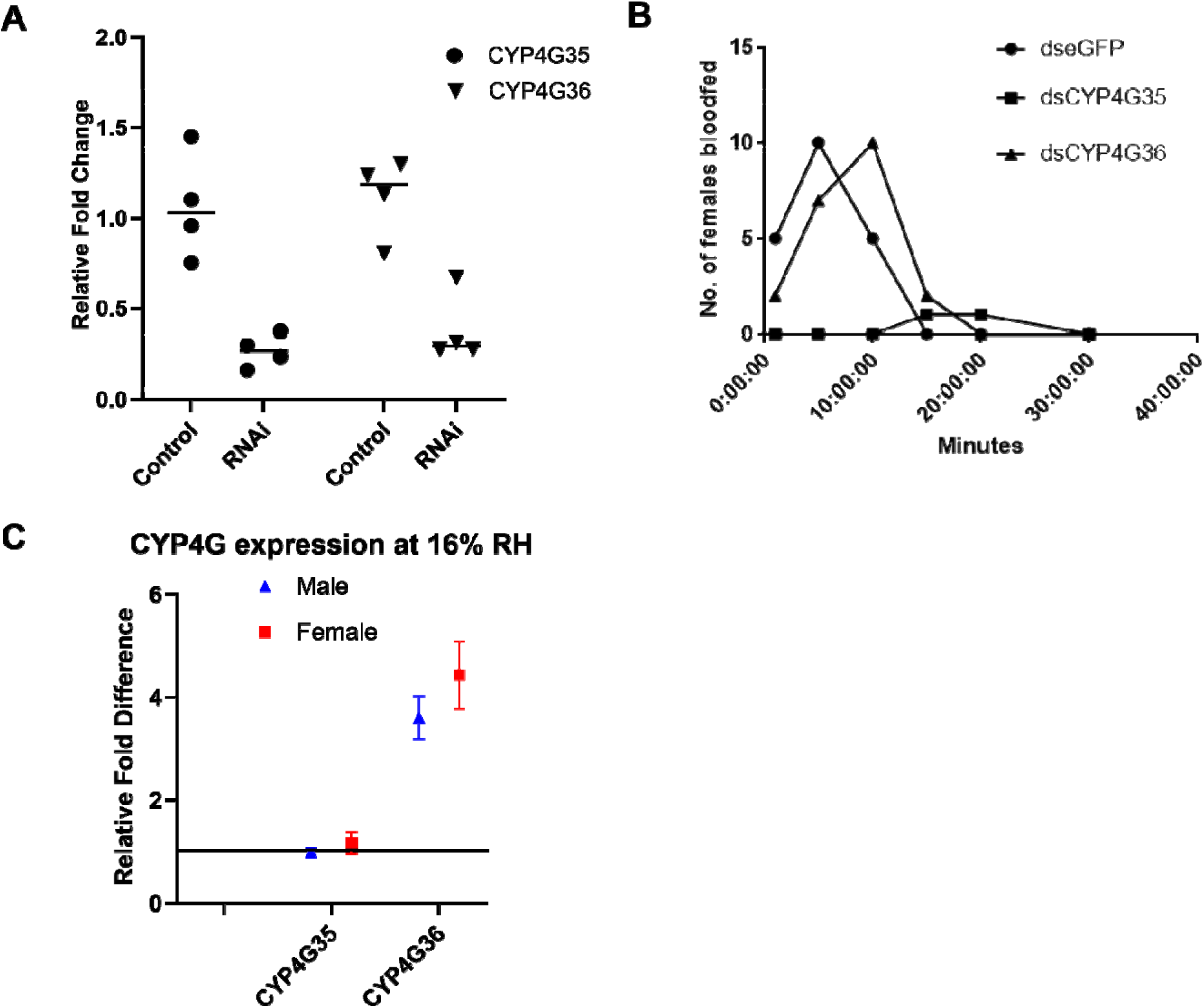
Effect of CYP4G knockdown on physiology and behavior. Transcript expression of CYP4G35 and CYP4G36 in adult females (A). Pupae were soaked in dsRNA solution, and RNA was extracted from adults eclosing from these pupae. An EGFP dsRNA was used as a control. n=4. Time taken by female mosquitoes from three different RNAi treatment groups to feed to repletion (B). The duration of blood feeding was recorded for each group and used to compare feeding efficiency across treatments. n=20. CYP4Gs expression at different humidity conditions (C). CYP4G35 and CYP4G36 knockdown male and female pupae were kept in an incubator at either 16% or 70% RH. 24h old adults eclosed from RNAi pupae were collected for RNA extraction and qRT-PCR. Mosquitoes reared at 70% were used as a control (solid horizontal line), and relative expression at 16% RH was determined.

Knockdown of either CYP4G35 or CYP4G36 had no adverse effect on the desiccation tolerance of adults, as all adults survived for up to 10 days post-RNAi at both 16 and 70% RH (Data not shown). The CYP4G36 transcript levels were higher at 16% RH (Fig. 6C), which might have led to increased CHC production; however, we did not test the CHC levels.

### CYP4G35 KO also confirms altered host seeking in an olfactometer assay

To explore the consequences of complete CYP4G knockout (KO), we generated CRISPR KO lines for both CYP4G35 and CYP4G36. Three single-guide RNAs (sgRNAs) were designed to target exon 1 of the CYP4G genes by manually identifying regions of the genome containing protospacer-adjacent motifs (PAMs) (Fig. S1, Table 2). In CYP4G35 mutants, a 4-base pair deletion was observed at the target site of one sgRNA in subsequent generations (Fig. 7A, Fig. S4). This deletion introduced a frameshift mutation (Fig. S5). Similarly, the sgRNA2 targeting CYP4G36 induced 5 bp deletions at its target site (Fig. 7A, Fig. S6), resulting in a frameshift mutation.

**Figure 7:**
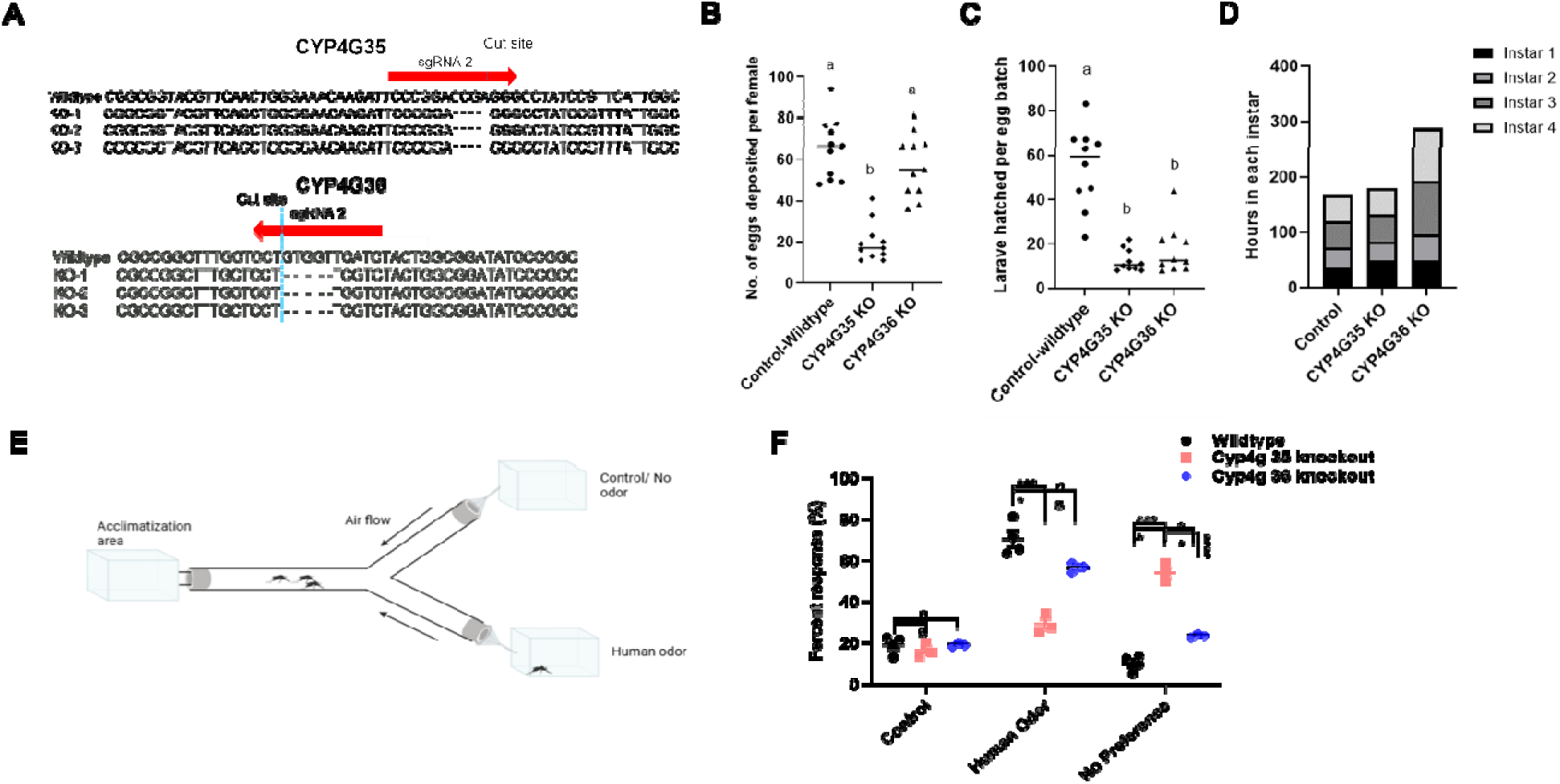
Knockout of CYP4G35 and CYP4G36 using CRISPR-Cas9. Embryos were either injected with three single-guide RNAs (multiplexed) targeting CYP4G35 or CYP4G36. Genomic DNA was extracted from the resulting G_0_ adults, and the sequences were determined by Sanger sequencing. Multiple alignments of the DNA sequences of the CYP4G35 (A) and CYP4G36 (B) of knockout (KO) versus the wild type (WT) illustrate deletions near the PAM site. A subset of G_2_ females was kept individually to count the number of eggs laid by CYP4G35 and CYP4G36 KO compared to the WT control (B). One-way ANOVA with Tukey’s multiple comparison was used (F (2, 30) = 36.91, P<0.0001). The number of larvae hatched from the eggs laid was counted (D). One-way ANOVA and Tukey’s comparison were used (F (2, 27) = 33.89, P<0.0001). Time spent in each larval instar until pupation was calculated (D). Schematic model of the Y-tube olfactometer used for a dual-choice attraction assay (E). Female mosquitoes were introduced into the long arm of the olfactometer and allowed to acclimate for 20 min before the assay began. A cotton sock worn for 10-12 h was used for human odor, and the no-odor-control received a similar sock that was not worn. Mosquitoes’ preference for human odor or control is plotted as a percent response (F). Statistical significance was assessed using one-way ANOVA and Dunnett’s multiple comparisons. ns = statistically nonsignificant *=p> 0.05, ** = p< 0.002, **** = p< 0.0001.

The CYP4G35 KO adults did not blood feed until at least 10d old, whereas WT adults fed as early as 2d post-emergence. Even at 10d post-eclosion, CYP4G35 KO females will only feed when the host is in direct contact (within the cage). The G_0_ hatch was much lower than expected, and only 14 individuals (10 females and 4 males) reached adulthood. None of the G_0_ females (outcrossed to WT) produced eggs even after a second blood meal. The WT females that were group mated to G_0_ males laid eggs; however, hatching was delayed. A subset of CYP4G35 KO adults was reared, blood-fed, and eggs laid and larval hatch were counted. While we did not weigh the mosquitoes to infer engorgement, only females with fully distended abdomens were used. The mean number of eggs laid and larvae hatched was significantly lower (∼70% reduction in eggs) (Fig. 7B, C) in CYP4G35 KO females compared to the WT females. Further, CYP4G35 eggs hatched after 2-3 days in water, whereas WT eggs in our insectary typically begin hatching within a few hours. Once hatched, the KO larvae developed similarly to WT larvae.

In CYP4G36 KO females, the number of eggs laid was similar to that in the control. However, ∼70% CYP4G36 KO eggs failed to hatch (Fig. 7B, C). The CYP4G36 KO larvae took longer to develop (on average, 4 days more than control or CYP4G35 KO) (Fig. 7D).

To evaluate host odor attraction in the CYP4G KO females, we employed a Y-tube olfactometer assay (Fig. 7E). Over 70% of CYP4G36 KO females oriented toward the human odor, closely aligning with the response of WT controls, where approximately 80% exhibited attraction (Fig. 7F). In contrast, only 20% of CYP4G35 KO females entered the odor arm, while the majority remained in the release zone and failed to respond to the stimulus (Fig. 7F).

### CYP4G KO affects the expression of other proteins and suggests compensatory mechanisms

A data-independent acquisition (DIA)- based global mass spectrometry proteomics approach was employed to identify proteins differentially expressed between CYP4G35 KO and WT samples. Principal Component Analysis (PCA) clearly separated the samples into four distinct groups: CYP4G35 KO head, KO body, WT head, and WT body, indicating distinct proteomic profiles (Fig. 8A). In total, 7,552 proteins were identified across all samples. Of these, 198 were differentially expressed in CYP4G35 KO mosquitoes, with 101 in the head and 96 in the body/carcass samples (Fig. 8B, C).

**Figure 8.**
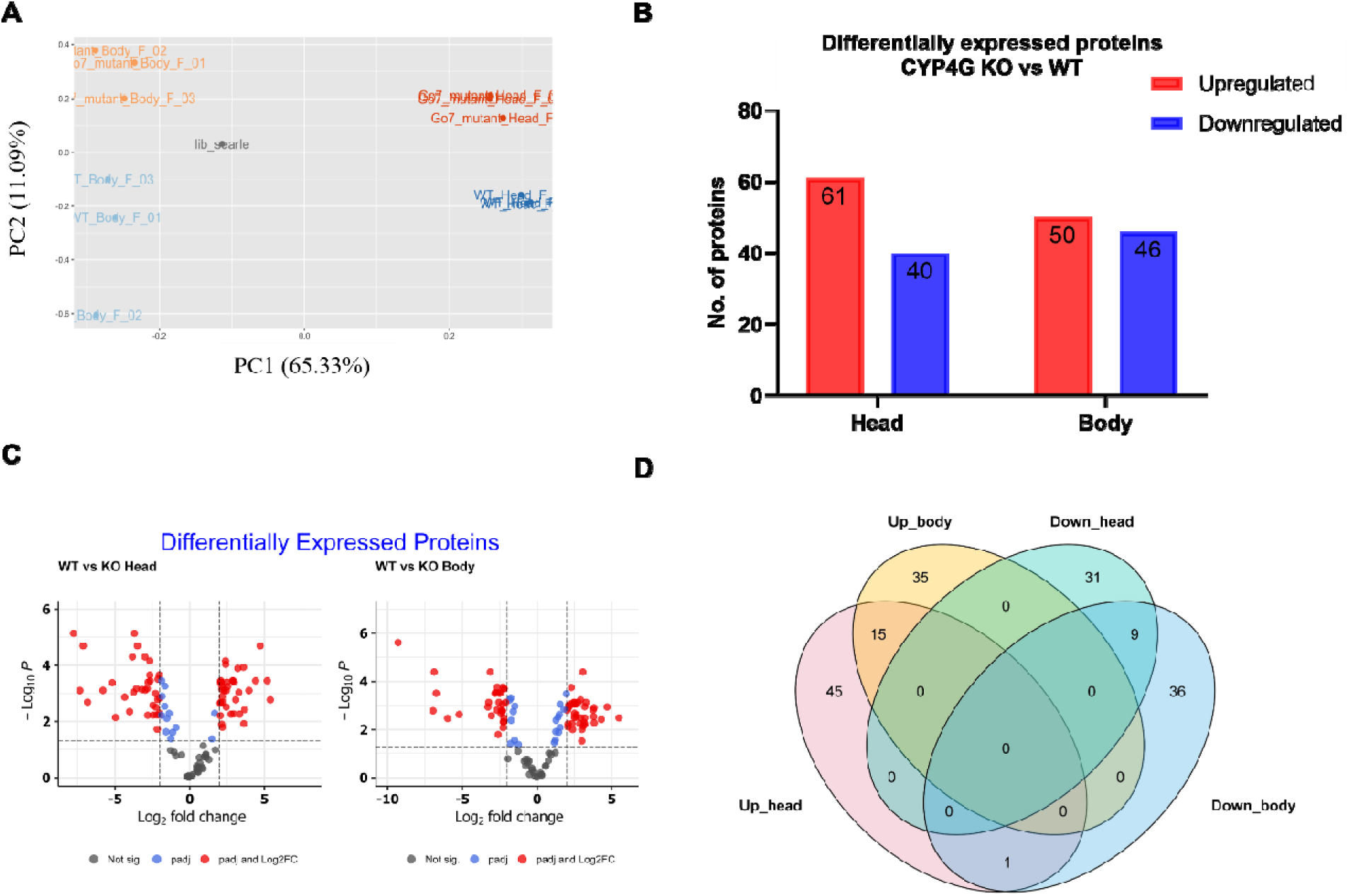
Proteomics. A) PCA plot illustrates variance stabilization normalization between data sets. The principal component analysis (PCA) shows the variance between the data set of head and body samples from both CYP4G mutant knock-out (CYP4G35 KO) and wild-type (WT). The red dots represent variance for head samples from CYP4G35 KO female *Ae. aegypti*. The orange dots represent variance for body samples from CYP4G35 KO female *Ae. aegypti*. The light blue dots represent the sample variance for body samples from wild-type female *Ae. aegypti*. The dark blue dots represent the sample variance for head samples from wild-type female *Ae. aegypti*. **B) Total differentially expressed proteins in head & body tissue in CYP4G35 KO vs. WT.** The number of DEPs up and down-regulated in head and body tissue in CYP4G35 KO. **C) Volcano plot of DEPs in carcass tissue of the CYP4G35 mutant vs. wild type**. Volcano plot showing the differentially expressed proteins in the body of *Ae. aegypti* between the CYP4G35 mutant versus wild type for 96 proteins included on all platforms. The red dots represent proteins that were upregulated and differentially expressed in the CYP4G35 knock-out mutant (Log fold-change >2, adjusted P<0.05). The blue dots represent proteins that were downregulated differentially expressed in the CYP4G35 knock-out mutant (Log fold-change >2, adjusted P<0.05). D) Venn diagram of up- and down-regulated proteins found in both the head and body of the CYP4G35 KO mutant.

Out of 101 differentially expressed proteins (DEPs) in the CYP4G35 KO heads, 40 were downregulated and 61 upregulated (Fig. 8B). 76 of the DEPs were exclusively in the head of the CYP4G35 KO. The rest were common between head and body samples (Fig. 8C). Of those upregulated in CYP4G35 KO heads, GO analysis indicated that the majority of proteins were associated with oxidoreductase activity, heme and iron binding, and microtubule binding (Fig. S7). Three CYPs: CYP6P12, CYP9J16, and CYP6A8, were upregulated in the head of the CYP4G35 KO females. The highest upregulated protein was a kinesin, AAEL004235 (kinesin motor domain) (Table 3, File S1), along with other microtubule-related proteins (KATNA1, AAEL013390). Two odor/pheromone binding proteins: OBP18 (AAEL004342) and OBP99c (AAEL011730) were also significantly upregulated (File S1).

KEGG pathway analysis suggested that retinol metabolism, bisphenol degradation, bile acid biosynthesis, methane metabolism, linoleic acid metabolism, biotin metabolism, and the biosynthesis pathways of cysteine, methionine, phenylalanine, tyrosine, and tryptophan were upregulated in CYP4G35 KO heads (Fig. S8). In general, proteases such as hydrolase, endopeptidase, and peptidase were downregulated. The downregulated KEGG pathways included insect hormone biosynthesis, chitin biosynthesis, steroid hormone, and the inositol pathway (Fig. S8).

In the CYP4G35 KO body (carcass) group, 96 DEPs were identified, 50 upregulated and 46 downregulated (Fig. 7B). 72 of 96 were exclusively in the CYP4G35 KO body (Fig. 8B, C). The KEGG pathway analysis of the upregulated proteins indicated a role in methane metabolism, arginine biosynthesis, and aminoacyl-tRNA biosynthesis (Fig. S8). The top upregulated proteins were AAEL026231 (pyridoxal phosphate-dependent aminotransferase), AAEL017010 (DNA-directed RNA polymerase subunit beta), AAEL026253 (ectopic P granules protein 5 homolog (EPG5)), AAEL024216 (carboxylesterase type B), and AAEL000832 (cleavage and polyadenylation specificity factor, CPSF) (Table 4, File S1). KEGG analysis of downregulated proteins in the body of the CYP4G35 KO mutants suggested a reduction in metabolic pathways: phenylpropanoid, alkaloid biosynthesis, lipopolysaccharide biosynthesis, glycerophospholipid metabolism, fructose/mannose, terpenoid biosynthesis, and inositol metabolism (Fig. S8, S9).

**Table 4.** Top five up or downregulated differentially expressed proteins in CYP4G35 KO body vs. WT body. DEPs from Gene Name, Gene ID, Fold change (Log2FC), and protein family the transcript belongs to.

| **Top 5 up or down-regulated DEPs** |  |  |  |
| --- | --- | --- | --- |
| **Gene Name** | **Gene ID** | **Log2FC** | **PFam Description** |
| **Upregulated in CYP4G35 KO Body vs. WT Body** |  |  |  |
| **Pyridoxal phosphate-dependent transferase** | AAEL026231 | 6.97 | Aminotransferase class-III; |
| **DNA-directed RNA polymerase** | AAEL017010 | 4.64 | DNA-directed RNA polymerase |
| **unspecified product** | AAEL026253 | 4.64 | Ectopic P granule protein 5 |
| **Carboxylesterase** | AAEL024216 | 4.6 | Carboxylesterase type B |
| **cleavage and polyadenylation specificity factor CPSF** | AAEL000832 | 4.24 | Cleavage/polyadenylation specificity factor |
| **Downregulated in CYP4G35 KO Body vs. WT Body** |  |  |  |
| **allantoinase** | AAEL007653 | -4.56 | Dihydroorotase |
| **unspecified product** | AAEL000950 | -5.09 | Low-temperature viability protein Ltv1 |
| **Nucleolar complex protein 2** | AAEL004818 | -6.15 | Nucleolar complex protein 2, Armadillo-type fold |
| **28 kDa heat- and acid-stable phosphoprotein (PDGF-associated protein), putative** | AAEL001952 | -7.04 | Casein kinase substrate, phosphoprotein PP28 |
| **Serine proteases, trypsin domain; Peptidase S1A** | AAEL009554 | -8.05 | chymotrypsin family; Peptidase S1, PA clan |

Of the 197 DEPs detected in either head or body, 15 proteins were upregulated in both body regions of CYP4G35 KO mosquitoes (Fig. 8D). Hemocyanins, specifically hexamerins (2-beta) (AAEL008045, AAEL008817, AAEL011169, and AAEL013757) were upregulated. Other more abundant proteins were AMP-dependent synthetase (AAEL002669), serine protease (AAEL011624), and serine/threonine protease (AAEL026028).

A total of nine proteins were downregulated in both head and body samples. These proteins consisted of dehydrogenases (AAEL026142, AAEL007669), hydrolases (AAEL010592, AAEL000642), and serine proteases (AAEL009554, AAEL024669, AAEL000252) (File S1).

Together, our data from multiple behavioral and biochemical assays are consistent with a role for CYP4G35 in modulating olfactory response. Based on the localization data that demonstrated CYP4G expression in antennal sensilla (base and shaft) and prior studies showing D. melanogaster CYP4G15 expression in glial cells, we propose a working model (Fig. 9) suggesting similar localization and function likely as ODE.

**Figure 9:**
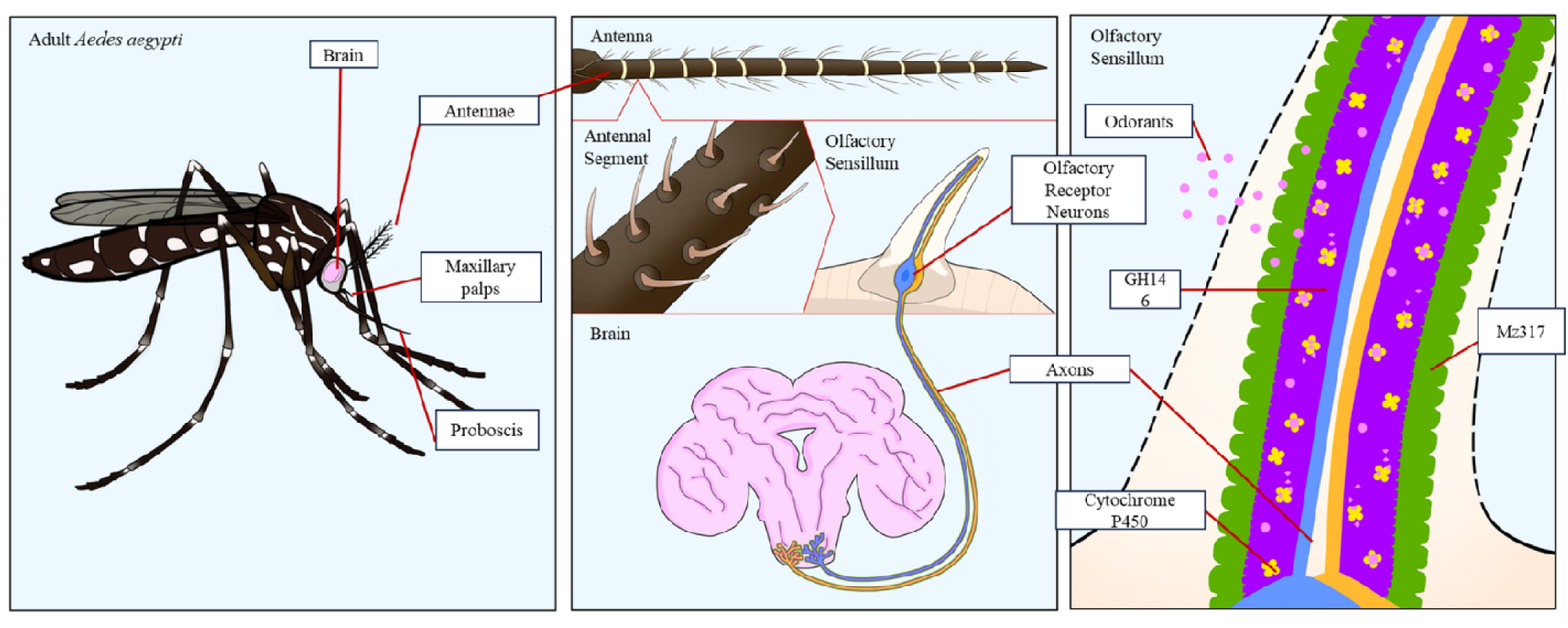
Working model of the CYP4G35 localization and function. Based on multiple behavioral and biochemical assays we propose a role for CYP4G35 in modulating olfactory response. Our localization data demonstrated CYP4G expression in antennal sensilla (base and shaft) and prior studies also confirmed *D. melanogaster* CYP4G15 expression in glial cells. We, therefore, propose a working model suggesting similar localization of CYP4G35 in GH146 glial cells and function likely as ODE.

## Discussion

The genomes of most Neoptera species carry at least two CYP4G genes, which are paralogs of the two characterized *D. melanogaster* CYP4G genes, CYP4G1 and CYP4G15. The duplication of the ancestral CYP4G is basal to Neoptera, and no CYP4G is found in Paleoptera, or beyond the class Insecta ^12^. All CYP4G sequences are characterized by a +44 residue insertion between the G and H helices ^12^. The *D. melanogaster* CYP4G1 and its orthologs in other insects are well-studied. These have an essential function of cuticular hydrocarbons biosynthesis for desiccation resistance, achieved mainly by the quantity of hydrocarbons produced, and to a lesser extent, in chemical communication by the blend of hydrocarbons produced ^12^. Similar to other dipterans, the *Ae. aegypti* genome expresses two CYP4Gs: CYP4G35 and CYP4G36. In the current study, transcript expression profiles across developmental stages showed the highest levels in late pupae before eclosion, when adult structures are forming. This expression profile aligns with the oxidative decarbonylase function of the well-studied orthologs of CYP4G36 (*D. melanogaster* CYP4G1 and *An. Gambiae* CYP4G16), as this enzyme is essential for cuticular hydrocarbon formation necessary for the incipient, terrestrial adult stage. The CYP4G transcripts were high in newly eclosed mosquitoes of both sexes, but the expression returned to the aquatic stage level once female blood-fed, likely due to higher water levels present in blood-fed mosquitoes. The expression likely remained consistent in older males and females after the blood meal was digested, but we did not extend our experiment beyond newly eclosed males and 24h post-blood-fed females.

CYP4G36 was expressed in all tissues examined, whereas CYP4G35 transcripts were most abundant in antennae (Fig. 1). This expression pattern agrees with the previously published neurotranscriptome of *Ae. aegypti* where CYP4G35 (AAEL006824) is highly expressed in female antennae compared to all other peripheral chemosensory tissues (including the brain, maxillary palp, proboscis, abdominal tip, and legs)^47^. A similar tissue-specific expression pattern was reported in other insects, where one or more CYP4G transcripts were abundant in olfactory tissues. For instance, in *A. mellifera*, CYP4G11 is highly expressed in antennae and legs and can accept short-chain aldehydes as substrates ^13^. These shorter-chain pheromonal and phytochemical alcohols and aldehydes, when metabolized by CYP4G11, yield volatile hydrocarbons. In the cabbage armyworm, *Mamestra brassicae*, CYP4G20 transcripts are abundantly expressed in the chemosensory tissues of adult males, suggesting a function in chemoreception ^22^. Similarly, in Asian citrus psyllid, *Diaphorina citri*, seven antennal-expressing CYP4G genes were identified, also suggesting a potential role in olfaction ^48^. Further, a study *in D. melanogaster* demonstrated that exposure to a plant odorant resulted in upregulation of several CYP genes in the antennae, reinforcing the idea that the CYPs might be contributing to the olfactory mechanism in insects ^49^.

RNAi-mediated reduction of CYP4G35 was associated with delayed host-orientation in an arm-in-cage assay. Similarly, the CYP4G35KO females failed to orient towards human odor and delayed feeding compared with WT controls. However, CYP4G36 KO females showed normal attraction to the human host and fed to engorgement. The KO phenotype in the Y-tube (Fig. 7H) isolates olfactory choice, whereas the RNAi arm-in-cage assay provides an ecologically relevant but multimodal readout because RNAi cohorts were not assayed in the Y-tube; therefore, the arm-in-cage delay reflects an integrated response to multiple cues. This pronounced behavioral deficit supports the conclusion that CYP4G35 plays a pivotal role in the detection or processing of host-related olfactory cues, whereas CYP4G36 appears dispensable for this function.

The CYP4G35 and CYP4G36 recombinant fusion proteins used for the substrate assay demonstrated that the CYP4G36 fusion protein was able to efficiently convert [9,10-³H_₂_]-octadecanol, to the corresponding hydrocarbon, thereby confirming its conserved enzymatic function with CYP4Gs characterized from other insects. However, CYP4G35 showed very little detectable activity toward heptadecanol, even after proper folding and expression, as confirmed by the CO and NADPH assays, suggesting a distinct physiological function. Long-chain aldehydes (≥C24) and a systematic chain-length panel were not tested here; thus, lack of CYP4G35 activity may reflect substrate scope.

The functional divergence of two CYP4Gs is further substantiated by our immunohistochemical and *in situ* hybridization analyses, which revealed that CYP4G35 is specifically localized within the sensillar lymph of the antennae and proboscis. The sensillar lymph is a specialized extracellular milieu where odorant-degrading enzymes (ODEs) facilitate the rapid inactivation and clearance of odorant molecules, thereby modulating olfactory signal dynamics ^6,50–52^. Within the antenna, auxiliary cells are thought to express ODEs, including CYPs, but few studies have examined their cellular localization. In mammals, ODEs have been detected in both olfactory neurons and surrounding support cells^49^. Mosquito CYPs may likewise be expressed in olfactory neurons in addition to their expected expression in auxiliary cells. This assumption is supported by the studies in *D. melanogaster*. CYP4G15 is expressed in the *Drosophila* brain ^23^, specifically in the cortex glial cell subtype ^16^. Both of these studies focused on the *Drosophila* brain; however, a recent transcriptomic study showed high expression of CYP4G15 in GH146 antennal glial cells ^53^. Antennae tissue in *D. melanogaster* is comprised of two primary glial cell populations: GH146 and Mz317 cells ^54^, wrapping glia and perineurial glia, respectively. About 30% of the glial cells in the antenna are GH146-type, which originate in the brain and migrate along the axons until they reach the third antennal segment, where they ensheath the axons of the olfactory sensory neurons (OSNs) that project from the antenna to the brain ^55,56^. Therefore, it is likely that the CYP4G15 associated with GH146 glial cells shows expression in both the brain and antennal tissue. Our IHC and ISH data show expression in *Ae. Aedes aegypti* antennal sensilla, where glial cells are present; however, we did not examine the brain because the expression in the head (brain included) was much lower than that of the antennae. While fluorescence was confined to the sensillar base/shaft, definitive assignment to extracellular lymph awaits co-labeling with lymph markers or ultrastructural localization.

Our proteomic analyses of the KO females provide additional insights into the systemic consequences of CYP4G35 deletion. In the head, CYP4G35 KO resulted in the upregulation of a kinesin and a microtubule-related protein (KATNA1). Two odor/pheromone binding proteins, OBP18 and OBP99c, were also significantly upregulated. Kinesin-1 is a heterotetrameric microtubule (MT) motor that plays diverse roles in both neuronal and non-neuronal cells, helping control the spatial and temporal organization of many cellular components through its ability to interact with various cargos ^57,58^. Both kinesin and KATNA1 (katanin) interact with microtubules but have opposing functions: kinesin transports cargo along microtubule tracks, while KATNA1 is an enzyme that severs or cleaves them. A conditional loss of kinesin in *D. melanogaster* reduces the electrophysiological response to odors and affects olfactory behavior. Orco, the olfactory receptor co-receptor, binds to the C-terminal tail fragments of the heterotrimeric kinesin motor, which is required to transfer Orco from the ciliary base to the outer segment of olfactory cilia ^59^. One hypothesis is that the disruption of odor degradation caused by the lack of CYP4G35 may result in increased transport of olfactory receptors and Orco to the cilia surface by kinesin to compensate for the change in olfactory behavior.

Another protein that was upregulated in both heads and body was ectopic P granule protein 5 (EPG5). The EPG5 protein is essential for the interaction between autophagosomes and lysosomes. The motility and positioning of lysosomes, driven by kinesin, are crucial for robust proteostasis ^60^. Similarly, two OBPs: OBP99c and OBP18, were upregulated. Both of these OBPs have been shown to express extracellularly in the head and antennae of *D. melanogaster* and the parasitoid wasp, *Microplitis mediator*, respectively ^61^. An intriguing finding was the upregulation of hexamerin in both the head and body of CYP4G35 KO females. Hexamerins (both Hex-1 and Hex-2) are stored in the fat body and the hemolymph of larvae and pupae ^62^. Earlier studies of newly eclosed female *Ae. aegypti* described numerous hexamerin protein granules in the fat body, which they suggested was remaining stored hexamerins; however, these granules disappeared within 3 days of eclosion ^63^. We collected head and body (carcass) samples from six-day-old females, and higher hexamerin in the KO samples suggests altered protein turnover. In addition, several oxidoreductases and cytochrome P450 enzymes, including CYP6P12, CYP9J16, and CYP6A8 were upregulated. CYP6A8 oxidizes lauric acid (C12:0), a short-chain unsaturated fatty acid, to produce 11-hydroxylauric acid, suggesting compensation for an enzyme that metabolizes short-chain carbon compounds ^64^. In contrast, the proteins downregulated in the head primarily included proteases and enzymes involved in cuticle remodeling and hormone biosynthesis pathways, such as those associated with chitin and steroid metabolism. This downregulation aligns with impaired cuticular barrier function and potential hormonal dysregulation, both of which are crucial for mosquito development, molting, and environmental resilience ^65–67^. Our finding that CYP4G35 KO negatively impacts egg laying and hatching also suggests hormonal dysregulation. Our data collectively suggest CYP4G35 as a specialized cytochrome P450 enzyme with a unique function in odor recognition, likely degradation, distinct from the canonical hydrocarbon-synthesizing CYP4Gs. This divergence highlights the functional versatility of P450 enzymes in mosquitoes and identifies CYP4G35 as a promising target for disrupting host-seeking behavior in vector control strategies.

While fluorescence is confined to the sensillar base and shaft, this pattern could originate from olfactory neuron dendrites, supporting cells, or cuticular interfaces rather than the extracellular lymph. Established methodologies using transmission electron microscopy with immunogold labeling have successfully achieved subcellular resolution to distinguish sensillar lymph from dendritic membranes ^68^, and co-immunolabeling with odorant-binding proteins has been used to confirm extracellular lymph localization ^69^ We plan to use these techniques. Additionally, we plan to explore CYP4G35 expression on antennal sections from mosquitoes in which subsets of olfactory neurons are labeled with GFP driven by transgenic reporters. In mosquitoes,□>□70% of olfactory neurons express Orco, therefore, Orco+ neurons may co-express CYP4G35. On the other hand, an IR-family co-receptor, Ir8a, is expressed by nearly all olfactory neurons that do not express Orco. Like Orco+ neurons, Ir8a+ neurons may express CYP4G35. Staining with special glial cell type markers will further help determine the localization of CYP4G35 in these cells.

## Data availability

Any additional information regarding the data reported in this paper is available from the LEAD Contact upon request.

## Authors’ contributions

M.G.-N., A.B.N., A.S., and D.M. designed the experiments. M.G.-N supervised the project. C.T. and G.B. provided technical support, troubleshooting, and intellectual contributions. A.S. and F.A. conducted the RNAi and relative gene expression assays, optimized the conditions, and validated these data. A.S., with the consultation from S.Y. and M.M., produced the fusion proteins for functional and localization studies. R.H. generated the knockout mosquitoes. O.G.-C. and M.P. carried out the *in-situ* hybridization and immunohistochemistry assays to determine the expression pattern of the proteins. J.P. analyzed the DIA proteomic data and helped with data interpretation and writing. M.G.-N. and A.S. wrote the first draft of the manuscript, and M.G.-N., A.B.N., A.S., and D.M. revised and finalized the text. All authors approved the final version of the manuscript.

## Acknowledgements

We would like to thank Dr. David Quilici and Ms. Rebekah J Woolsey of the Mick Hitchcock, Ph.D. Nevada Proteomics Center (RRID:SCR 017761), UNR, for proteomics data acquisition. We would also like to thank <u>Channa Aluvihare</u>, Insect Transformation Center, for rearing the CRISPR KO lines up to the second generation. This work was supported by the NIH Neuroscience COBRE Award (P20GM103650) to MG-N and the National Institute of General Medical Sciences (GM103440) from the National Institutes of Health.

