## Supplemental materials for "A cytochrome P450 G subfamily member, CYP4G35, is highly expressed in antennae and modulates olfactory response in *Aedes aegypti* mosquitoes"

**Supplementary Materials**

**Figure Legends:**

**Figure S1:** Gene maps showing exons (black boxes) and introns (lines) and designed sgRNAs targeting the first exon.

**Figure S2:** Sequencing used for making fusion proteins CYP4G35-CPR and CYP4G36-CPR

**Figure S3: CYP4G35 Antibody validation by Dot-Blot:** showed the reaction of Dot-Blot assay. By Dot-Blot assay, mosquito specific CYP4G35 antibody (1:500, 1:1000, and 1:5000) was detected in triplicate from extracted mosquito proteins, no antibody as negative control and selected peptides as positive control.

**Figure S4: CYP4G35 knockout in *Ae. Aegypti* females**. **A)** Representative figure showing the sequences of CYP4G35 knockout females (G0) aligned to wildtype (WT), **B)** Editing efficiency as access by using ICE synthego CRISPR analysis tool (https://www.synthego.com/products/bioinformatics/analysis).

**Figure S5:** Amino acid translations for CYP4G35 KO mutants resulting in a truncated protein compared to the wild type.

**Figure S6: CYP4G36 knockout in *Ae. Aegypti* females**. **A)** Representative figure showing the sequences of CYP4G35 knockout females (G0) aligned to wildtype (WT), **B)** Editing efficiency as access by using ICE synthego CRISPR analysis tool ((https://www.synthego.com/products/bioinformatics/analysis).

**Figure S7. Bubble plot of molecular function enrichment for CYP4G35 KO head tissue.** GO classification of biological process and molecular function for proteins upregulated in head CYP4G35 KO versus WT.

**Figure S8. KEGG analysis of the metabolic pathways within CYP4G35 KO and WT.** The metabolic pathway was identified to be upregulated and downregulated in the body proteome for CYP4G35 KO and WT. The metabolic pathway for identified to be upregulated and downregulated in the head proteome for CYP4G35 KO and WT.

**Figure S9. Bubble plot of molecular function enrichment for CYP4G35 KO body tissue.** GO classification of biological process and molecular function for proteins upregulated in body CYP4G35 KO versus WT (left).

**Figure S1**

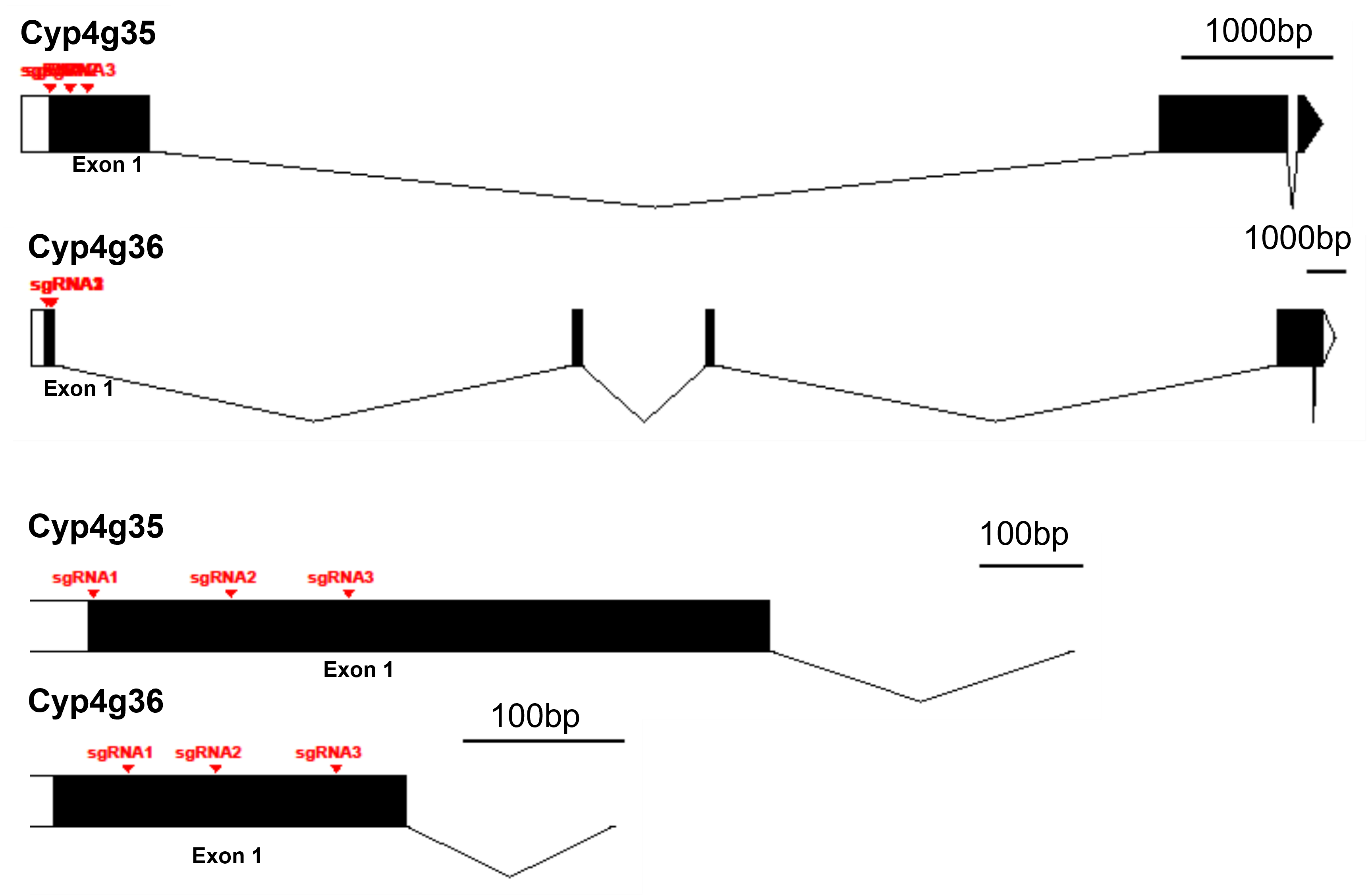

**Figure S2**

**For CYP4G35:**

To clone into pENTR4(Nco-) amplified with pENTR4F4/R5:

R5:	CCGGATCCAGTCGACTGAAT

(R5ic):	ATTCAGTCGACTGGATCCGG

To create a fusion-clone with CPR, amplify pENTR4_4G2/CPR with pENTR4R5 and CPRFus_F2

CPRFus_F2:	tcaactacatctatacaaccc

CPRFus_F2ic	GGGTTGTATAGATGTAGTTGA

CYYP4G35_orf:

>XM_001659099.2 Aedes aegypti AAEL008345-RA (CYP4G35), partial mRNA

ATGAGCGCGGAAATTGTGGCCGAAAGGGGCAGCAGCCTGGTGTCGCTGGCCGTGCCGATGGTCATTTTTATGACCCTGGTCTTGGTAGCCAGTGCATTGTTCCATTTCTGGATGATATCCCGGCGGTACGTTCAACTGGGAAACAAGATTCCCGGACCGAGGGCCTATCCGTTCATTGGCAATGCCAATATGCTGCTGGGGATGAACCACAATGAGATCATGGAGCGGGCGATGCAGTTGAGTTATATCTATGGAAGTGTGGCTCGGGGCTGGCTCGGGTATCATTTGGTGGTGTTTTTGACCGAGCCAGCGGACATTGAGATCATCCTGAACAGTTATGTGCACCTGACAAAGTCCAGCGAGTACAGGTTTTTCAAGCCATGGCTTGGCGATGGGCTGTTGATCAGCAGTGGCGAAAAGTGGCGATCACATCGGAAGCTAATCGCTCCGGCGTTCCATATGAATGTCCTGAAGACGTTCGTGGATGTGTTCAACGATAACAGTTTGGCGGTGGTGGAACGGATGCGAAAGGAGGTGGGGAAGGAGTTCGACGTGCACGACTATATGAGTGAAGTTACGGTGGATATTCTGCTGGAGACGGCCATGGGATCGCAGAGGACGAGCGAGAGCAAGGAGGGATTTGATTATGCGATGGCTGTGATGAAAATGTGTGACATCCTACACTCCCGTCAGTTAAAATTCCACCTTCGGATGGACTCCGTCTTCAACTTCACCAAAATCAAGCAGGAACAGGAACGTCTGCTCGGCATCATCCACGGCCTCACCCGAAAAGTCGTCAAACAGAAGAAGGAACTCTTCGAGAAGAATTTTGCCGACGGAAAGCTGCCTTCGCCGTCCCTTTCCGAAATTATTGCCAAGGAAGAGTCCGAATCCAAAGAACCGCTTCCGGTCATCTCGCAGGGTTCGCTCCTCAGGGACGATCTGGACTTCAACGATGAGAATGACATCGGCGAGAAGCGAAGGCTTGCCTTCCTGGACCTGATGATCGAAACGGCCAAGAGCGGTGCCGATCTGACCGATGAAGAGATCAAAGAAGAAGTCGACACCATCATGTTTGAAGGACACGACACCACTGCGGCTGGATCCAGCTTTGTGCTGTGCCTTCTCGGCATTCACCAGGACGTTCAAGATCGAGTTTACAAAGAAATCTACCAGATCTTTGGCAACTCCAAGCGGAAAGCTACATTCAACGACACCTTGGAGATGAAGTACCTGGAACGGGTGATCTTTGAAACCCTGAGAATGTATCCACCGGTTCCGGTGATTGCCCGTAAAGTTACACAAGATGTCCGGCTGGCATCCCACGACTACGTGGTTCCAGCCGGAACCACGGTCGTCATCGGAACTTATAAAGTGCACCGACGAGCGGACATTTACCCTAATCCAGATGTGTTCAACCCGGACAATTTCCTACCGGAACGCACACAGAATCGCCACTACTACAGCTACATCCCATTCAGCGCCGGACCGCGAAGTTGCGTCGGTAGAAAATATGCCATGCTGAAACTAAAGGTCCTTCTGTCAACCATCCTGCGCAACTACAGGGTCGTGTCAAATCTCAAGGAATCGGACTTTAAGCTACAAGGCGACATTATCCTGAAACGGACCGATGGCTTCAGAATACAGCTGGAACCGAGAGTCTAA

>XP_001659149.1 AAEL008345-PA [Aedes aegypti]

MSAEIVAERGSSLVSLAVPMVIFMTLVLVASALFHFWMISRRYVQLGNKIPGPRAYPFIGNANMLLGMNHNEIMERAMQLSYIYGSVARGWLGYHLVVFLTEPADIEIILNSYVHLTKSSEYRFFKPWLGDGLLISSGEKWRSHRKLIAPAFHMNVLKTFVDVFNDNSLAVVERMRKEVGKEFDVHDYMSEVTVDILLETAMGSQRTSESKEGFDYAMAVMKMCDILHSRQLKFHLRMDSVFNFTKIKQEQERLLGIIHGLTRKVVKQKKELFEKNFADGKLPSPSLSEIIAKEESESKEPLPVISQGSLLRDDLDFNDENDIGEKRRLAFLDLMIETAKSGADLTDEEIKEEVDTIMFEGHDTTAAGSSFVLCLLGIHQDVQDRVYKEIYQIFGNSKRKATFNDTLEMKYLERVIFETLRMYPPVPVIARKVTQDVRLASHDYVVPAGTTVVIGTYKVHRRADIYPNPDVFNPDNFLPERTQNRHYYSYIPFSAGPRSCVGRKYAMLKLKVLLSTILRNYRVVSNLKESDFKLQGDIILKRTDGFRIQLEPRV

>CYP4G35 Sf9 bias

attcagtcgactggatccggATGTCCGCTGAGATCGTGGCTGAGCGCGGTTCCTCCCTGGTGTCCCTGGCTGTGCCTATGGTGATCTTCATGACCCTGGTGCTGGTGGCTTCCGCTCTGTTCCACTTCTGGATGATCTCCCGCCGCTACGTGCAGCTGGGTAACAAGATCCCTGGTCCTCGCGCTTACCCTTTCATCGGTAACGCTAACATGCTGCTGGGTATGAACCACAACGAGATCATGGAGCGCGCTATGCAGCTGTCCTACATCTACGGTTCCGTGGCTCGCGGTTGGCTGGGTTACCACCTGGTGGTGTTCCTGACCGAGCCTGCTGACATCGAGATCATCCTGAACTCCTACGTGCACCTGACCAAGTCCTCCGAGTACCGCTTCTTCAAGCCTTGGCTGGGTGACGGTCTGCTGATCTCCTCCGGTGAGAAGTGGCGCTCCCACCGCAAGCTGATCGCTCCTGCTTTCCACATGAACGTGCTGAAGACCTTCGTGGACGTGTTCAACGACAACTCCCTGGCTGTGGTGGAGCGCATGCGCAAGGAGGTGGGTAAGGAGTTCGACGTGCACGACTACATGTCCGAGGTGACCGTGGACATCCTGCTGGAGACCGCTATGGGTTCCCAGCGCACCTCCGAGTCCAAGGAGGGTTTCGACTACGCTATGGCTGTGATGAAGATGTGCGACATCCTGCACTCCCGCCAGCTGAAGTTCCACCTGCGCATGGACTCCGTGTTCAACTTCACCAAGATCAAGCAGGAGCAGGAGCGCCTGCTGGGTATCATCCACGGTCTGACCCGCAAGGTGGTGAAGCAGAAGAAGGAGCTGTTCGAGAAGAACTTCGCTGACGGTAAGCTGCCTTCCCCTTCCCTGTCCGAGATCATCGCTAAGGAGGAGTCCGAGTCCAAGGAGCCTCTGCCTGTGATCTCCCAGGGTTCCCTGCTGCGCGACGACCTGGACTTCAACGACGAGAACGACATCGGTGAGAAGCGCCGCCTGGCTTTCCTGGACCTGATGATCGAGACCGCTAAGTCCGGTGCTGACCTGACCGACGAGGAGATCAAGGAGGAGGTGGACACCATCATGTTCGAGGGTCACGACACCACCGCTGCTGGTTCCTCCTTCGTGCTGTGCCTGCTGGGTATCCACCAGGACGTGCAGGACCGCGTGTACAAGGAGATCTACCAGATCTTCGGTAACTCCAAGCGCAAGGCTACCTTCAACGACACCCTGGAGATGAAGTACCTGGAGCGCGTGATCTTCGAGACCCTGCGCATGTACCCTCCTGTGCCTGTGATCGCTCGCAAGGTGACCCAGGACGTGCGCCTGGCTTCCCACGACTACGTGGTGCCTGCTGGTACCACCGTGGTGATCGGTACCTACAAGGTGCACCGCCGCGCTGACATCTACCCTAACCCTGACGTGTTCAACCCTGACAACTTCCTGCCTGAGCGCACCCAGAACCGCCACTACTACTCCTACATCCCTTTCTCCGCTGGTCCTCGCTCCTGCGTGGGTCGCAAGTACGCTATGCTGAAGCTGAAGGTGCTGCTGTCCACCATCCTGCGCAACTACCGCGTGGTGTCCAACCTGAAGGAGTCCGACTTCAAGCTGCAGGGTGACATCATCCTGAAGCGCACCGACGGTTTCCGCATCCAGCTGGAGCCTCGCGTGtcaactacatctatacaaccc

Amplify this insert with R5ic/CPRFusF2ic

pENTR4_R5ic	ATTCAGTCGACTGGATCCGG

CPRFusF2ic	GGGTTGTATAGATGTAGTTGA

Mix with vector/CPR fragment (ampmlified with R5/CPRFusF2) for Gibson assembly.

**For CYP4G36:**

To clone into pENTR4(Nco-) amplified with pENTR4F4/R5:

R5:	CCGGATCCAGTCGACTGAAT

(R5ic):	ATTCAGTCGACTGGATCCGG

To create a fusion-clone with CPR, amplify pENTR4_4G2/CPR with pENTR4R5 and CPRFus_F2

CPRFus_F2:	tcaactacatctatacaaccc

CPRFusF2ic	GGGTTGTATAGATGTAGTTGA

CYP4G36

>XM_001648326.2 Aedes aegypti AAEL004054-RA (CYP4G36), partial mRNA

ATGTCCGCGACGGTTGCCCCAGCGGACCCCGTGATGGCGAATGCCAACATCGCGTCGCCCATGAATGTGTTTTACTTCCTGCTGGCGCCGGCTTTGCTCCTGTGGTTCATCTACTGGCGGATATCCCGGCAACATATGCTGAAGCTGGCTGAGAAAATACCTGGCCCACCTGGATTACCCCTGCTTGGAAACGCGCTGGAACTGATTGGAACCTCTCATTCCGTCTTCCGCAACGTAATCGAGAAAGGAAAGGACTTCAATCAGGTCATCAAAATCTGGATTGGACCGAAGCTGATCGTCTTCCTGGTGGATCCACGTGATGTAGAACTATTGCTCAGCAGTCACGTGTACATCGACAAATCTCCGGAATATCGCTTCTTCAAGCCTTGGCTGGGAAATGGACTGCTCATCAGTACAGGTCACAAATGGCGTCAACATCGCAAACTCATTGCTCCCACTTTCCATTTGAACGTGCTCAAGAGCTTCATCGATTTGTTCAACGAGAACTCCCGGCTGGTGGTGGAAAAGATGCACAAGGAGGCCGGCAAAACCTTCGATTGCCATGACTACATGAGCGAGTGTACCGTGGAAATCCTGCTCGAAACCGCCATGGGAGTTTCGAAGAAAACTCAGGACCAATCTGGATTCGACTATGCTATGGCTGTAATGAAGATGTGCGACATCCTGCATCTGCGTCATCGTAAGATGTGGCTCTACCCGGACCTCTTCTTCAACATGTCCCAGTATGCCAAGCGTCAGGTTAAACTCCTGGACACCATCCACAGCTTGACCCGCAAGGTCATCCGTAACAAGAAGGCCGCCTTTGCTACCGGAACACGAGGATCGCTGGCCACAACTTCCATCAAGACGGCTGAATTTGAAAAACCGAAATCCAACATCAACACCAATAGCGTGGAAGGACTATCTTTTGGTCAGTCGGCTAATCTGAAGGATGATTTGGACGTCGATGAAAATGATGTCGGAGAGAAGAAGCGATTGGCATTCCTTGATCTGCTGTTAGAAAGCGCAGAAAACGGTGCTTTGATCTCGGATGAAGAAATCAAAAATCAAGTCGACACTATTATGTTTGAAGGACACGACACGACTGCAGCCGGAAGTAGCTTCTTCCTTTCGATGATGGGCATCCACCAGCACATCCAAGACAAAGTCATCCAAGAATTGGATGACATTTTCGGAGACTCGGATCGACCTGCTACATTCCAGGATACTTTAGAGATGAAATATCTTGAGCGATGCTTGATGGAAACGCTGCGAATGTATCCTCCAGTACCAATTATTGCAAGATCTCTGAAACAGGACTTGAAGCTGGCATCCAGCGATTTAGTTGTACCATCTGGCGCCACTATCGTCGTCGCTACATACAAATTGCATCGACTTGA

AACAATCTACCCGAATCCCAATGTGTTTGATCCCGACAACTTCCTCCCTGAAAGACAGGCCAACCGCCACTATTATGCATTCGTTCCATTCTCAGCCGGACCCAGAAGTTGTGTTGGTCGCAAGTACGCTATGCTCAAGCTGAAGGTCATCCTTTCCACCATTTTGCGAAACTTCCGCGTCATCTCGGACCTCAAAGAAGAAGACTTCAAACTACAGGCTGATATCATTCTGAAACGAGAAGAAGGCTTCCAGATCCGCCTGGAGCCCCGCCAACGTAAACCAAAAGCAGCGAAAGCCTGA

CYP4G36

>XP_001648376.1 AAEL004054-PA [Aedes aegypti]

MSATVAPADPVMANANIASPMNVFYFLLAPALLLWFIYWRISRQHMLKLAEKIPGPPGLPLLGNALELIGTSHSVFRNVIEKGKDFNQVIKIWIGPKLIVFLVDPRDVELLLSSHVYIDKSPEYRFFKPWLGNGLLISTGHKWRQHRKLIAPTFHLNVLKSFIDLFNENSRLVVEKMHKEAGKTFDCHDYMSECTVEILLETAMGVSKKTQDQSGFDYAMAVMKMCDILHLRHRKMWLYPDLFFNMSQYAKRQVKLLDTIHSLTRKVIRNKKAAFATGTRGSLATTSIKTAEFEKPKSNINTNSVEGLSFGQSANLKDDLDVDENDVGEKKRLAFLDLLLESAENGALISDEEIKNQVDTIMFEGHDTTAAGSSFFLSMMGIHQHIQDKVIQELDDIFGDSDRPATFQDTLEMKYLERCLMETLRMYPPVPIIARSLKQDLKLASSDLVVPSGATIVVATYKLHRLETIYPNPNVFDPDNFLPERQANRHYYAFVPFSAGPRSCVGRKYAMLKLKVILSTILRNFRVISDLKEEDFKLQADIILKREEGFQIRLEPRQRKPKAAKA

>CYP4G36_bias

ATTCAGTCGACTGGATCCGGATGTCCGCTACCGTGGCTCCTGCTGACCCTGTGATGGCTAACGCTAACATCGCTTCCCCTATGAACGTGTTCTACTTCCTGCTGGCTCCTGCTCTGCTGCTGTGGTTCATCTACTGGCGCATCTCCCGCCAGCACATGCTGAAGCTGGCTGAGAAGATCCCTGGTCCTCCTGGTCTGCCTCTGCTGGGTAACGCTCTGGAGCTGATCGGTACCTCCCACTCCGTGTTCCGCAACGTGATCGAGAAGGGTAAGGACTTCAACCAGGTGATCAAGATCTGGATCGGTCCTAAGCTGATCGTGTTCCTGGTGGACCCTCGCGACGTGGAGCTGCTGCTGTCCTCCCACGTGTACATCGACAAGTCCCCTGAGTACCGCTTCTTCAAGCCTTGGCTGGGTAACGGTCTGCTGATCTCCACCGGTCACAAGTGGCGCCAGCACCGCAAGCTGATCGCTCCTACCTTCCACCTGAACGTGCTGAAGTCCTTCATCGACCTGTTCAACGAGAACTCCCGCCTGGTGGTGGAGAAGATGCACAAGGAGGCTGGTAAGACCTTCGACTGCCACGACTACATGTCCGAGTGCACCGTGGAGATCCTGCTGGAGACCGCTATGGGTGTGTCCAAGAAGACCCAGGACCAGTCCGGTTTCGACTACGCTATGGCTGTGATGAAGATGTGCGACATCCTGCACCTGCGCCACCGCAAGATGTGGCTGTACCCTGACCTGTTCTTCAACATGTCCCAGTACGCTAAGCGCCAGGTGAAGCTGCTGGACACCATCCACTCCCTGACCCGCAAGGTGATCCGCAACAAGAAGGCTGCTTTCGCTACCGGTACCCGCGGTTCCCTGGCTACCACCTCCATCAAGACCGCTGAGTTCGAGAAGCCTAAGTCCAACATCAACACCAACTCCGTGGAGGGTCTGTCCTTCGGTCAGTCCGCTAACCTGAAGGACGACCTGGACGTGGACGAGAACGACGTGGGTGAGAAGAAGCGCCTGGCTTTCCTGGACCTGCTGCTGGAGTCCGCTGAGAACGGTGCTCTGATCTCCGACGAGGAGATCAAGAACCAGGTGGACACCATCATGTTCGAGGGTCACGACACCACCGCTGCTGGTTCCTCCTTCTTCCTGTCCATGATGGGTATCCACCAGCACATCCAGGACAAGGTGATCCAGGAGCTGGACGACATCTTCGGTGACTCCGACCGCCCTGCTACCTTCCAGGACACCCTGGAGATGAAGTACCTGGAGCGCTGCCTGATGGAGACCCTGCGCATGTACCCTCCTGTGCCTATCATCGCTCGCTCCCTGAAGCAGGACCTGAAGCTGGCTTCCTCCGACCTGGTGGTGCCTTCCGGTGCTACCATCGTGGTGGCTACCTACAAGCTGCACCGCCTGGAGACCATCTACCCTAACCCTAACGTGTTCGACCCTGACAACTTCCTGCCTGAGCGCCAGGCTAACCGCCACTACTACGCTTTCGTGCCTTTCTCCGCTGGTCCTCGCTCCTGCGTGGGTCGCAAGTACGCTATGCTGAAGCTGAAGGTGATCCTGTCCACCATCCTGCGCAACTTCCGCGTGATCTCCGACCTGAAGGAGGAGGACTTCAAGCTGCAGGCTGACATCATCCTGAAGCGCGAGGAGGGTTTCCAGATCCGCCTGGAGCCTCGCCAGCGCAAGCCTAAGGCTGCTAAGGCTtcaactacatctatacaaccc

Amplify this insert with R5ic/CPRFusF2ic

pENTR4_R5ic	ATTCAGTCGACTGGATCCGG

CPRFusF2ic	GGGTTGTATAGATGTAGTTGA

Mix with vector/CPR fragment (amplified with R5/CPRFusF2) for Gibson assembly.

**Figure S3**

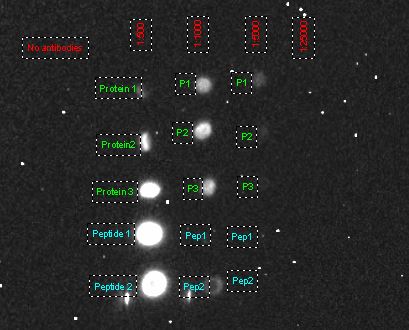

**Figure S4**

**A)**

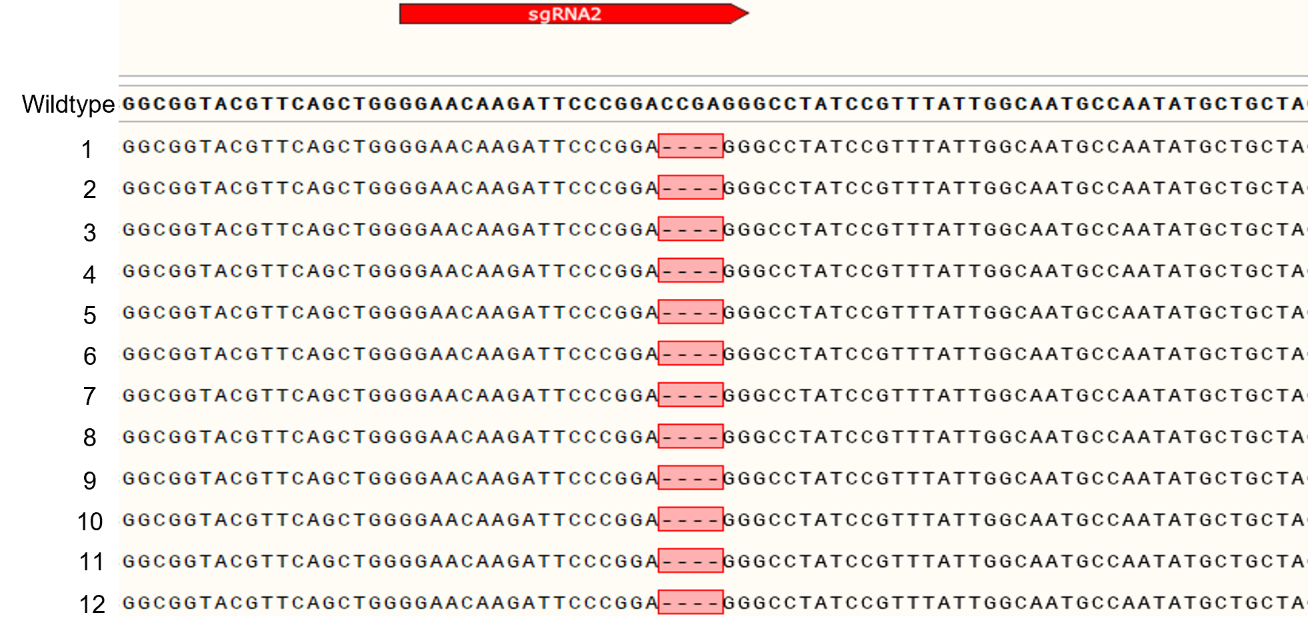

**B)**

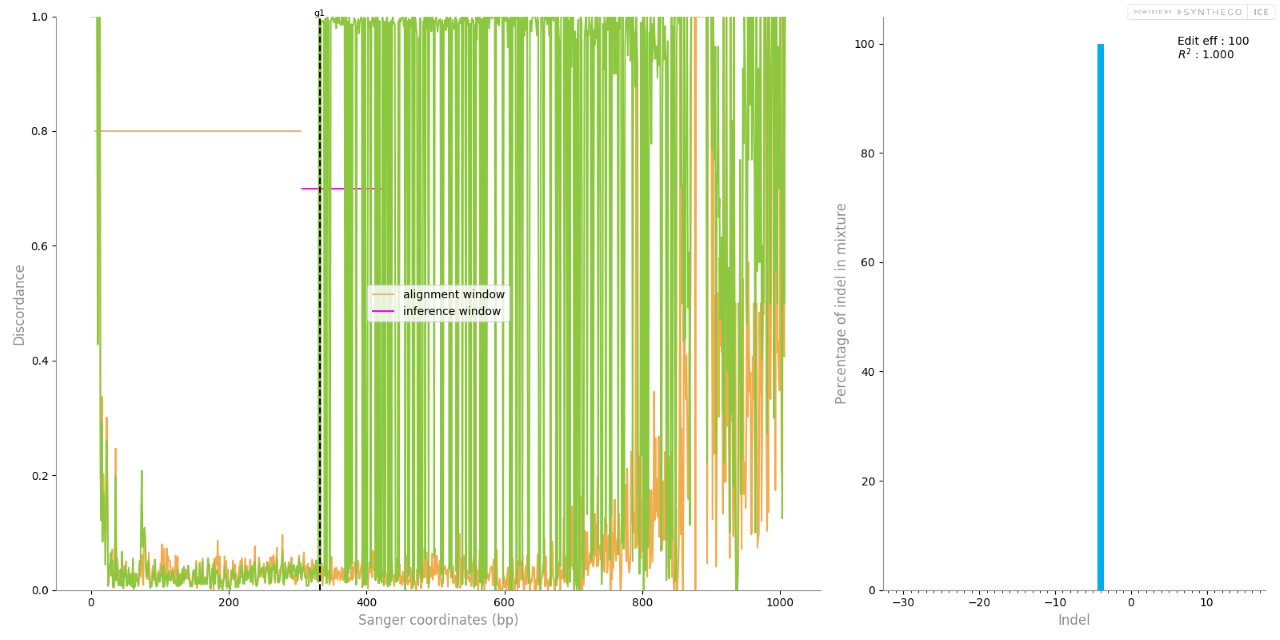

**Figure S5:**

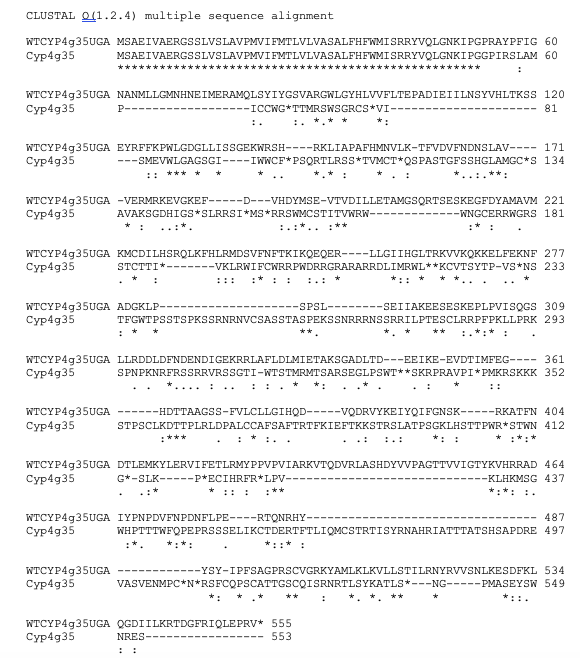

**Figure S6**

**A)**

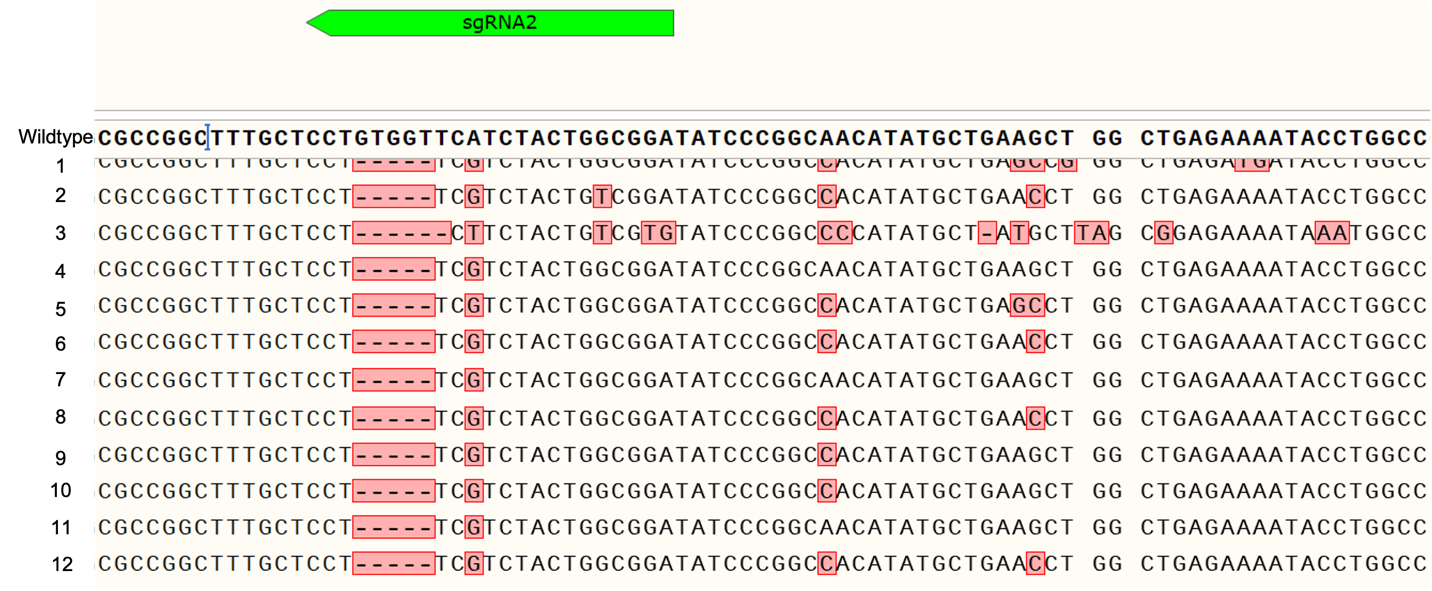

**B)**

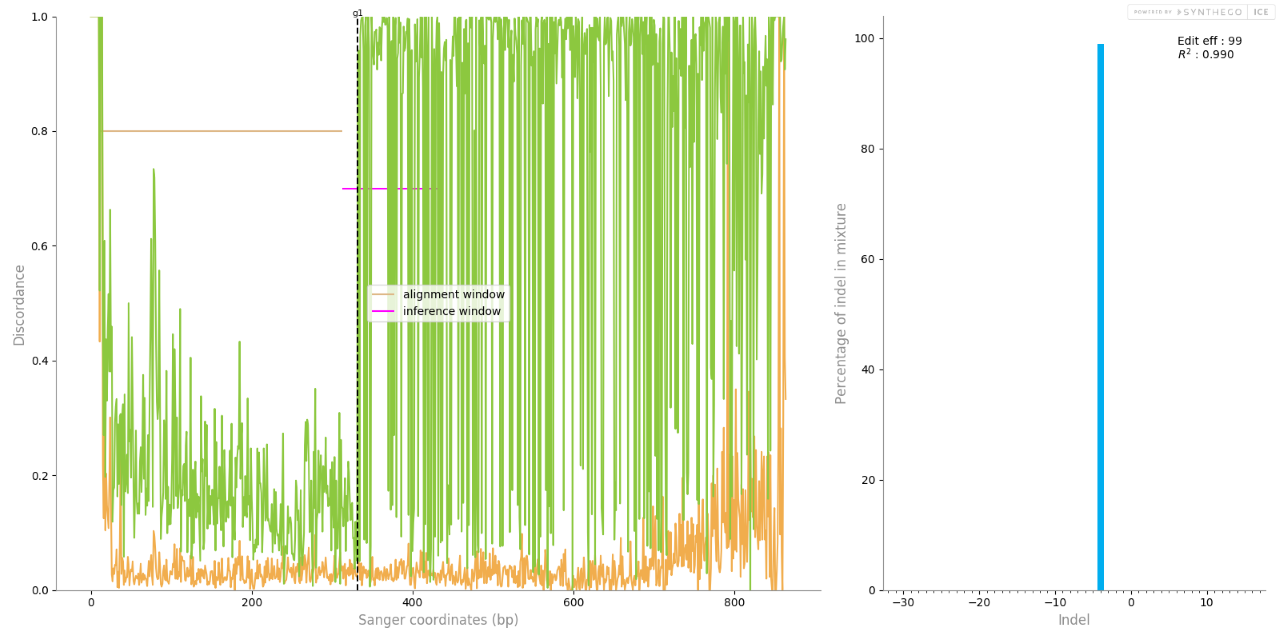

**Figure S7**

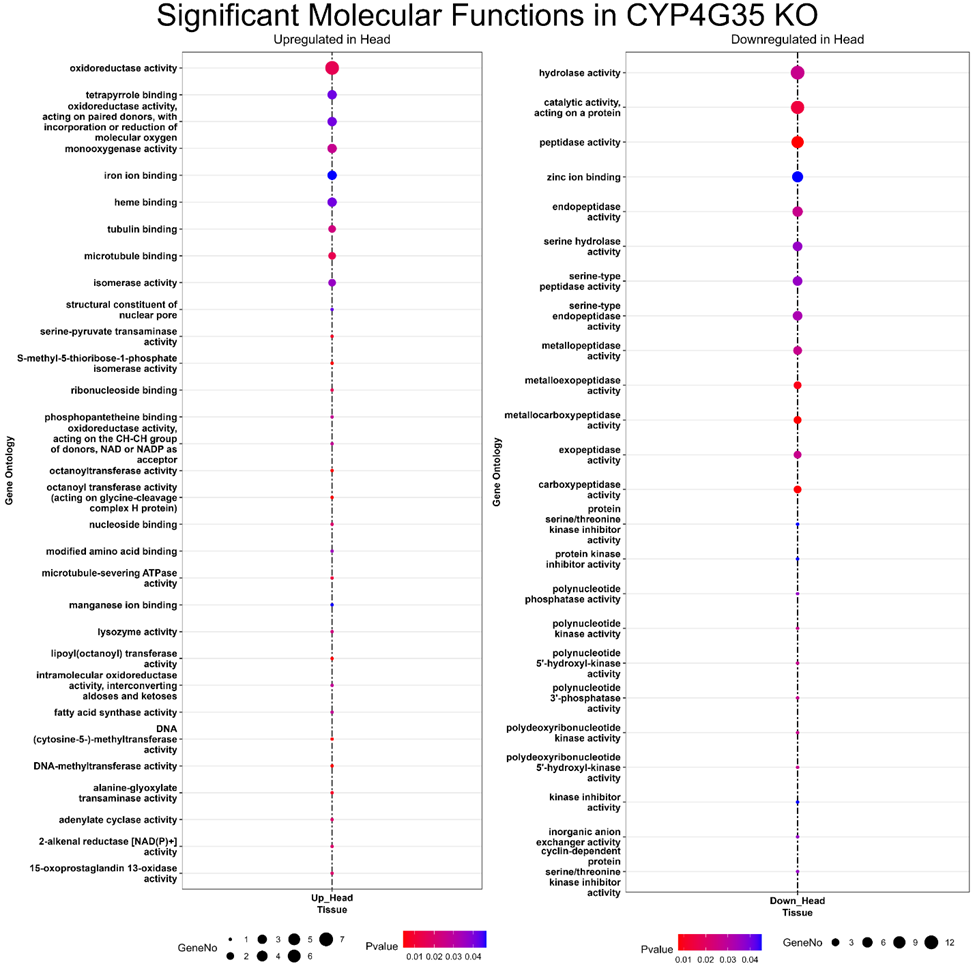

**Figure S8**

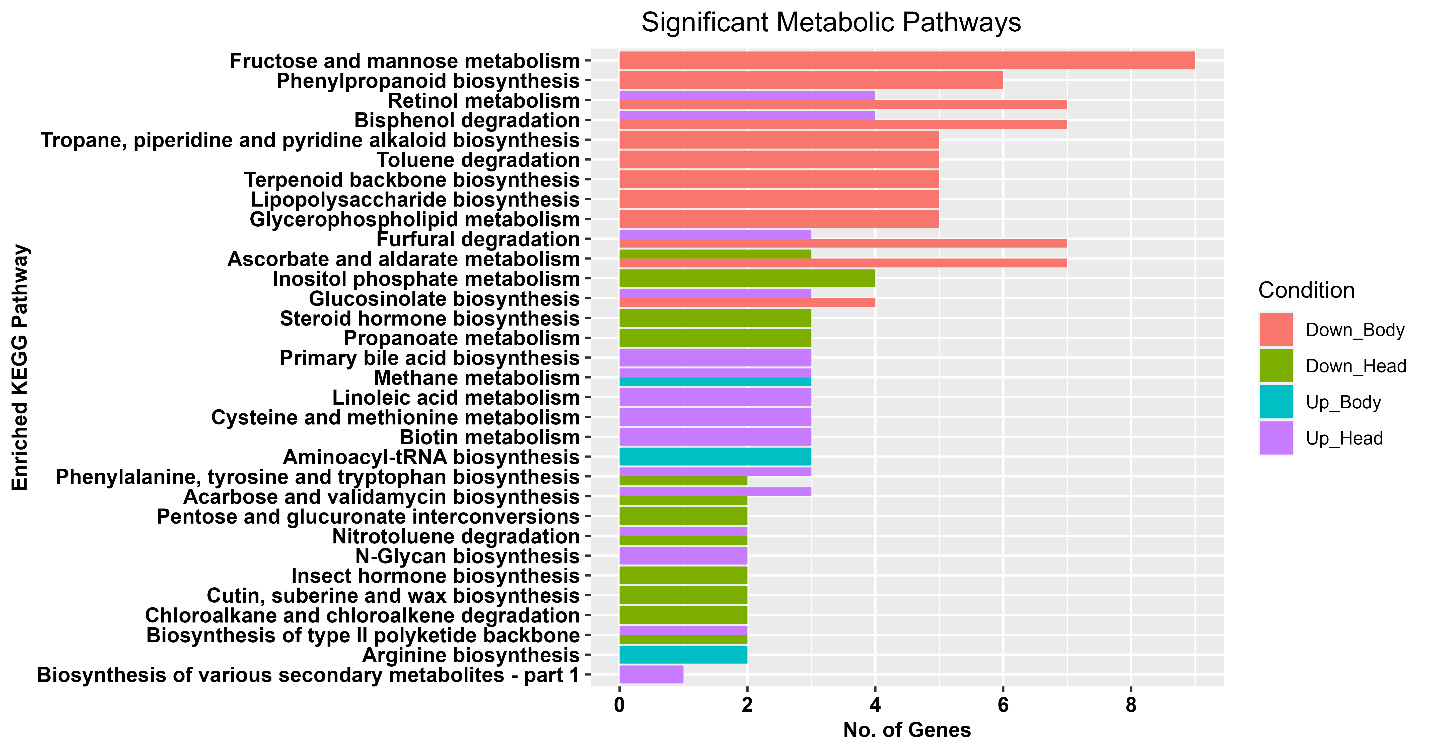

**Figure S9**

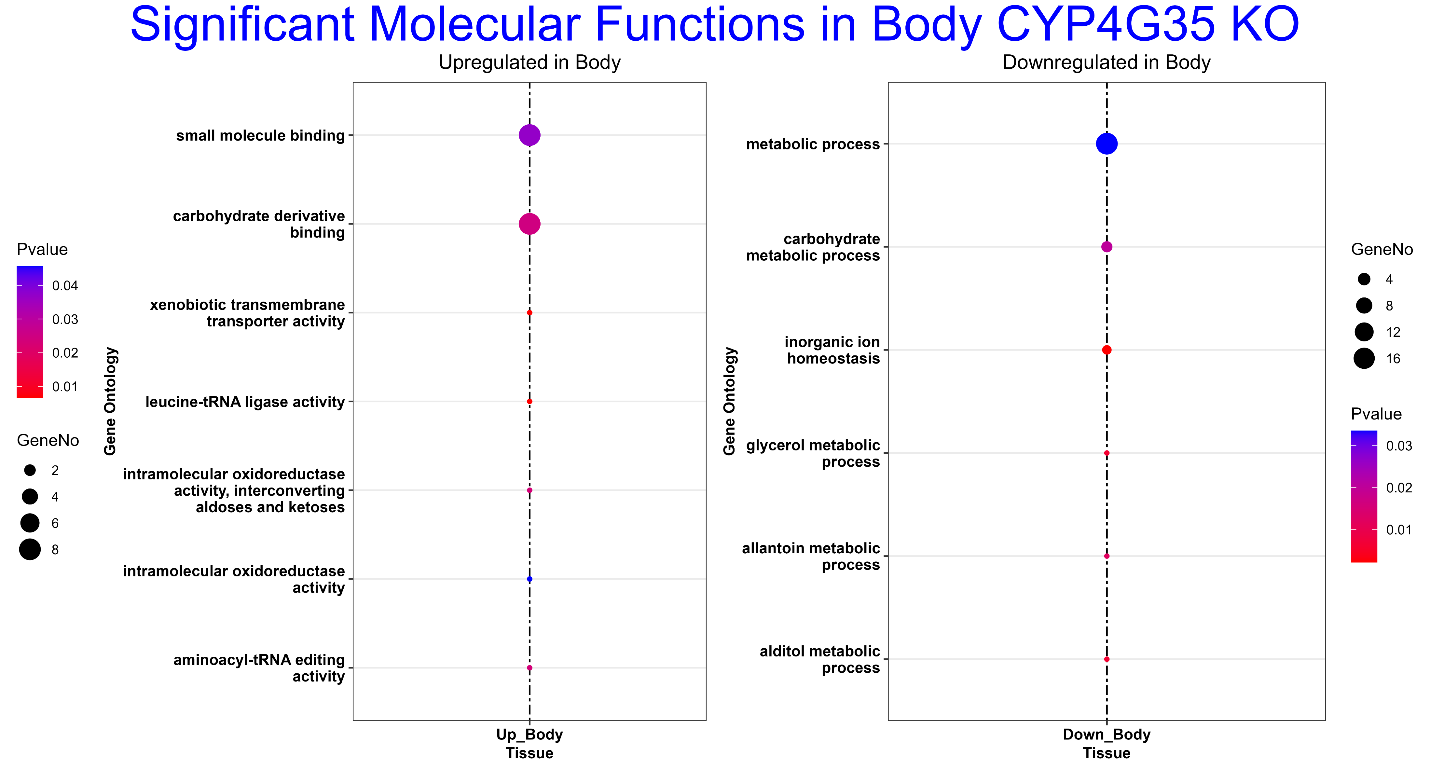
